# Protein detection in blood with single-molecule imaging

**DOI:** 10.1101/2021.05.19.444873

**Authors:** Chih-Ping Mao, Shih-Chin Wang, Yu-Pin Su, Ssu-Hsueh Tseng, Liangmei He, Annie A. Wu, Richard B.S. Roden, Jie Xiao, Chien-Fu Hung

## Abstract

The ability to identify and characterize individual biomarker protein molecules in patient blood samples could enable diagnosis of diseases at an earlier stage, when treatment is typically more effective. Single-molecule imaging offers a promising approach to accomplish this goal. However, thus far single-molecule imaging methods have only been used to monitor protein molecules in solutions or cell lysates, and have not been translated into the clinical arena. Furthermore, the detection limit of these methods has been confined to the picomolar (10^−12^ M) range. In many diseases, the circulating concentrations of biomarker proteins fall several orders of magnitude below this range. Here we describe Single-Molecule Augmented Capture (SMAC), a single-molecule imaging technique to visualize, quantify, and characterize individual protein molecules of interest down to the subfemtomolar (<10^−15^ M) range, even in complex biologic fluids. We demonstrate SMAC in a wide variety of applications with human blood samples, including the analysis of disease-associated secreted proteins, membrane proteins, and rare intracellular proteins. Using ovarian cancer as a model, a lethal malignancy in which high-grade disease is driven almost universally by alterations in the *TP53* gene and frequently only diagnosed at a late, incurable stage, we found that mutant pattern p53 proteins are released into the blood in patients at an early stage in disease progression. SMAC opens the door to the application of single-molecule imaging in non-invasive disease profiling and allows for the analysis of circulating mutant proteins as a new class of highly specific disease biomarkers. The SMAC platform can be adapted to multiplex or high-throughput formats to characterize heterogeneous biochemical and structural features of circulating proteins-of-interest.

**One Sentence Summary:** A single-molecule imaging approach detects individual disease-specific protein molecules, including mutant intracellular proteins, in blood samples.

## Main Text

Diseased cells release biomarker proteins into the bloodstream (*1*). These proteins can be tissue-specific but normal in structure, contain mutated regions, or carry abnormal secondary modifications. Conventional blood tests, such as the enzyme-linked immunosorbent assay (ELISA), typically cannot discern protein concentrations below the picomolar (pM; 10^−12^ M) range (*1*). The circulating levels of biomarker proteins associated with early stages of common disorders such as cancer or infection frequently fall in the femtomolar (fM; 10^−15^ M) range and below (*2, 3*). Newer methods, including digital ELISA (*4*), DNA biobarcoding (*5*), proximity ligation (*6, 7*), and immuno-PCR (*8, 9*), have been developed to improve the sensitivity of protein assays to the fM range. However, these tests rely on enzymatic amplification and ensemble measurements of the target molecule (*1*). Ensemble methods are limited by detection errors from background or non-specific reagent binding, especially in complex clinical fluids such as blood.

Single-molecule imaging approaches visualize individual protein molecules, providing greater sensitivity, reliability, and depth-of-information than ensemble methods (*10*). The detection limit of single-molecule imaging approaches has thus far reached the pM range in cell lysates (*11, 12*). However, single-molecule imaging of proteins in the blood has not been previously achieved. Here we describe Single-Molecule Augmented Capture (SMAC), a technique that allows quantification of individual protein molecules in the blood down to the sub-fM range. SMAC offers orders of magnitude greater detection sensitivity and specificity than other single-molecule imaging methods, opening up the possibility of, for the first time, quantifying and characterizing disease-associated molecules in patient samples at the single-molecule level. SMAC interrogates images formed by individual target protein molecules within biological samples and uses a fluorescence shape recognition algorithm to correct detection errors derived from non-specific antibody absorption or autofluorescence in complex biological fluids. Thus, true signals are reliably distinguished from false background signals in samples. Here we demonstrate a wide variety of applications of SMAC in blood-based human disease profiling.

In SMAC, individual proteins-of-interest are continuously pulled down by a capture antibody on a microfluidic device, probed by a fluorophore-labeled detection antibody, and visualized by single-molecule imaging (Fig. 1A). We achieved sub-fM sensitivity by implementing the following strategies (Fig. S1A). First, we created the SMAC chip, a highly efficient target-capture microfluidic device (Fig. 1A). The chip has the following features: (a) it is coated it with a dense layer of multi-valent, biotinylated antibody via a NeutrAvidin linker (*12, 13*), which enhances capture affinity and suppresses non-specific binding; (b) the total capture area of the chip is minimized, which concentrates proteins-of-interest by >10^4^-fold; and (c) a staggered herringbone micromixer roof (*14*) and oscillating sample flow scheme are incorporated onto the chip, which promote target-antibody collisions (Fig. S1A-C). Second, we employed a flow cytometry-based antibody screening process to rapidly identify the best capture/detection antibody pairs for target proteins (Fig. S1D, E). Third, we conducted time-stream averaged total internal reflection fluorescence (TIRF) microscopy images to resolve spatially individual fluorescent spots of target protein molecules (Fig. S2A, B; see Methods for details). Time-stream TIRF microscopy helps overcome diffusive background from autofluorescent substances in test samples and in the microfluidic device itself. These autofluorescent substances dissociate rapidly from the chip surface and photobleach more quickly than fluorophore-labeled detection antibodies. By contrast, detection antibodies specifically bound to target protein molecules remain attached to the chip for a longer time (Fig. S2C). Thus, by time-averaging an imaging stream, SMAC removes autofluorescent background signals while preserving specific signals from detection antibodies (Fig. S2A, B).

**Fig. 1.**
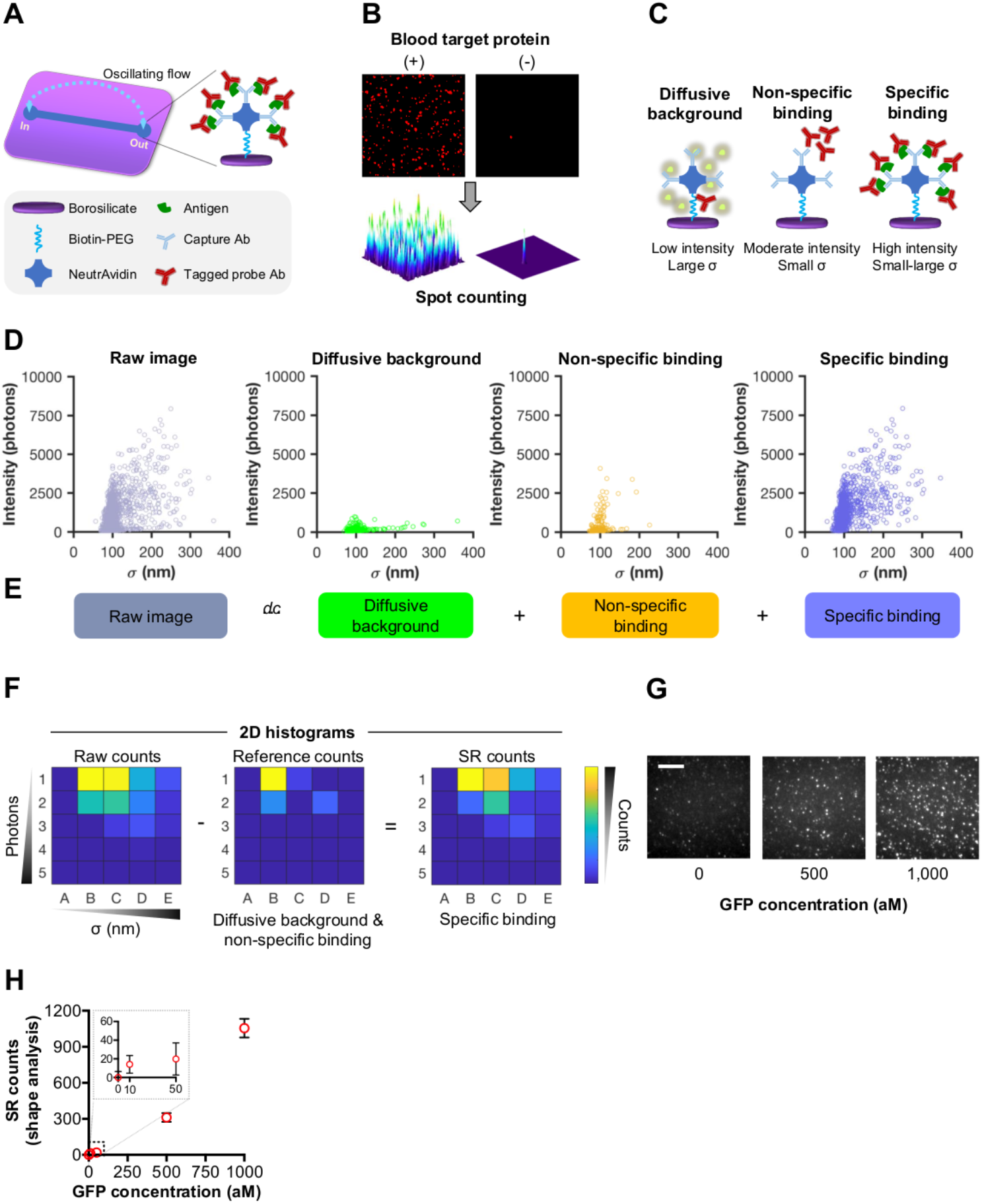
SMAC and protein analysis methods. (A) Schematic diagram of the SMAC platform. Proteins-of-interest were pulled down as clusters via continuous oscillating flow on a multivalent microfluidic device and then probed with fluorophore-labeled detection antibody. (B) Target protein clusters were visualized by TIRF microscopy. (C) Schematic diagrams depicting different binding types that give rise to different fluorescence intensity and spot size combinations. Scatter plots (D) and decomposition (E) of spot sizes (σ) and intensities arising from different binding types after Gaussian fitting of each spot. These data were converted into a two-dimensional (2D) histogram of intensity and σ as shown in (F). (F) The number of specific binding spots (SR counts) is obtained by subtracting the 2D histogram of a scaled reference histogram conveying the intensity-σ distributions of diffusive background and non-specific binding from the 2D histogram of raw counts (see Methods for details). (G) Representative SMAC images of purified GFP molecules at 500 aM and 1 fM concentrations. The intrinsic fluorescence of GFP was measured without detection antibody. (H) Graph illustrating the sensitivity of SMAC with shape analysis (SR counts) using purified GFP from 10 aM to 1 fM. Data are expressed as mean ± SD. Scale bar, 4 μm.

Finally, to achieve sub-fM sensitivity with minimal detection errors for proteins of rare occurance in samples, we applied a ‘fluorescence shape’ recognition algorithm (referred to as shape analysis; see Methods for details). Because NeutrAvidin and the capture antibody are multi-valent (Fig. 1A), fluorophore-labeled detection antibody molecules form clusters around target protein molecules on each NeutrAvidin tetramer, generating fluorescent spots with combinations of size (measured by the *σ* of the Gaussian fitting of the spot) and intensity (*I*; measured in number of photons) that are distinct from those of background spots due to diffusive background and non-specifically absorbed detection antibody molecules (Fig. 1B, C). We referred to these combinations of size and intensity as the *I-σ* shape (Fig. 1D). The *I-σ* shape reflects the combination of fluorescent signals emitted by specific antibody binding, non-specific antibody binding, and background diffusive autofluorescence and can be deconvoluted into its individual components via shape analysis (Fig. 1D, E). To perform shape analysis, we first represented the *I-σ* shape of a test sample as a two-dimensional (2D) histogram depicting the absolute number of spots in a set number of *I-σ* bins (Fig. 1F). We corrected for detection errors by subtracting out the maximum projected number of background reference spots (experimentally derived from a large pool of negative control background samples) from each bin of the test sample *I-σ* histogram (Fig. 1F). This analysis allowed us to overcome confounding effects of background signals at extremely low target protein concentrations in complex fluids such as blood.

We first validated the design of SMAC using a capture antibody targeting purified GFP. Because GFP is intrinsically fluorescent, we did not use a detection antibody. By applying the shape recognition algorithm, we achieved a limit of detection (LOD) of 61 aM GFP (Fig. 1G, H and Fig. S3A). This detection sensitivity is >10^4^-fold more sensitive than ELISA and existing single-molecule imaging approaches (*11, 12*) (Fig. S3B). Note that for samples containing proteins of relatively high abundance, such as >10 fM GFP, it is not necessary to use the shape-recognition algorithm. Instead, we measured the overall fluorescence intensity per sample (referred to as integrated intensity analysis) by adjusting the EMCCD (electron-multiplier charge-coupled device) camera gain such that the signal would fall within the linear range of the camera. We then compensated the image signal level using the corresponding EM gain and a standard calibration curve (see Methods for details), allowing us to accurately detect GFP from sub-fM concentrations up to 100 nM in a single sample without the need for dilution (Fig. S3A), which corresponds to a dynamic range of around nine orders of magnitude (compared to approximately two orders of magnitude for ELISA).

Next, we developed SMAC to detect a disease-associated secreted protein, prostate-specific antigen (PSA), as a proof of principle for the application of SMAC to clinical samples (Fig. 2A). PSA is a well-established biomarker for prostate cancer (*15–18*). While the normal prostate gland constitutively produces PSA, the expression of this protein is often dysregulated in prostate cancer cells, leading to either elevated or reduced PSA levels in the blood (*15–18*). SMAC could detect purified PSA at an LOD of 648 aM (Fig. 2B, C), 10^5^ times below the limit of the current clinical PSA test (*4*) (Fig. S4B) and is comparable to the reported limit of digital ELISA (*4*). SMAC was able to detect PSA in lysate derived from a single prostate cancer cell titrated into aqueous buffer or human blood (Fig. 2D-F). Furthermore, the SMAC PSA assay achieved a dynamic range of over six orders of magnitude (from 5 fM to >1 nM) (Fig. S4A) compared to two orders of magnitude for the clinical PSA test.

**Fig. 2.**
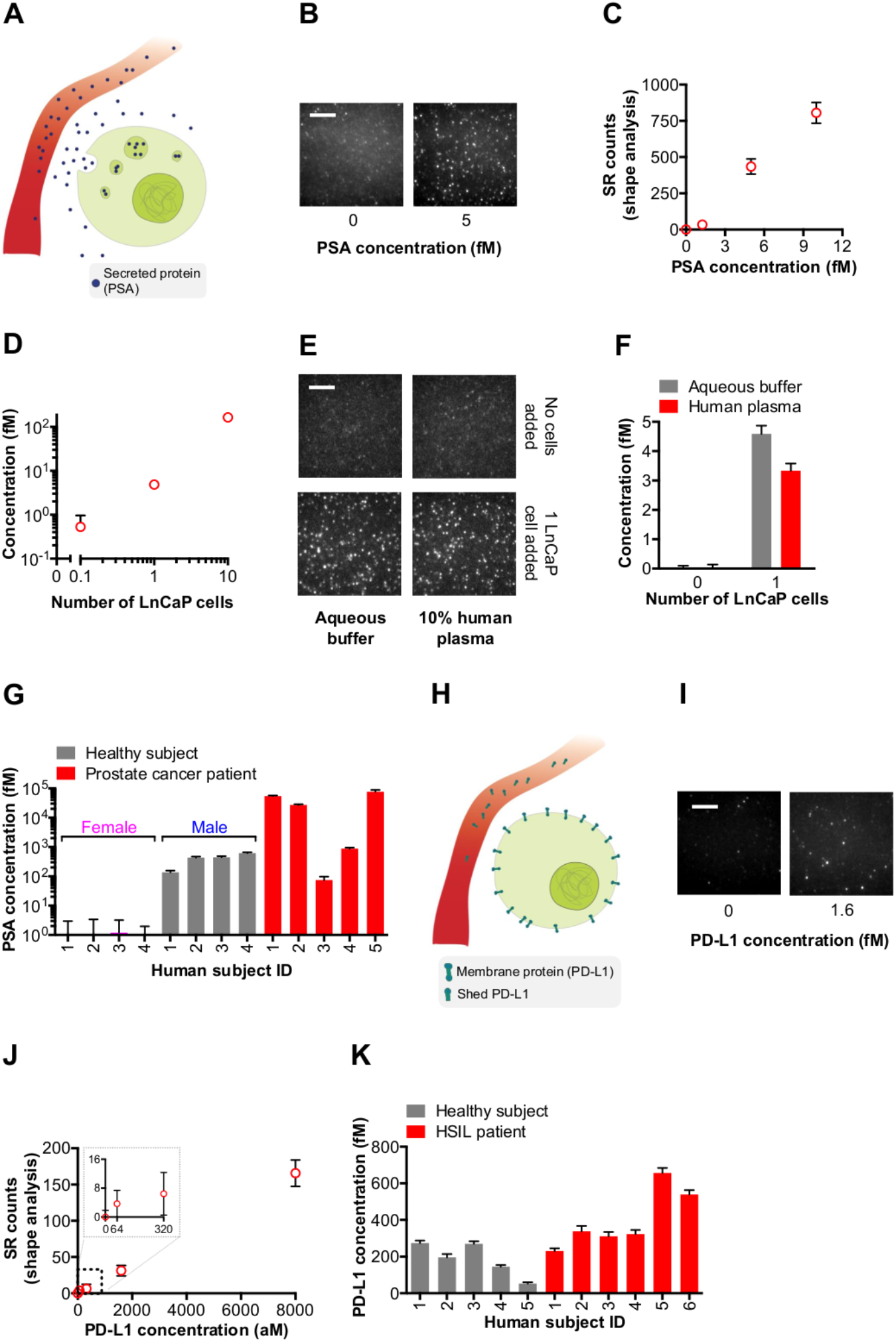
Detection of secreted and membrane proteins in blood by single-molecule imaging. (A) Schematic diagram of secreted PSA release from a tumor cell (lime) into a blood vessel (red). SMAC images (B) and shape analysis (C) of purified human PSA at fM concentrations in aqueous buffer. (D) Quantification of PSA in lysate from different numbers of human prostate cancer cells (LnCaP) added into aqueous buffer. SMAC images (E) and quantification of PSA (F) in lysate from one LnCaP cell in either aqueous buffer or human plasma. (G) PSA levels in the blood of prostate cancer patients (*n* = 5) and healthy male (*n* = 4) and female (*n* = 4) control blood donors. (H) Schematic diagram of membrane-bound PD-L1 release from a tumor cell (lime) into a blood vessel (red). SMAC images (I) and shape analysis (J) of purified human PD-L1 at fM concentrations in aqueous buffer. (K) Quantification of circulating PD-L1 levels in patients with high-grade squamous intraepithelial lesions (HSIL; *n* = 6) and healthy donors (*n* = 5). PSA data and PD-L1 data are expressed as mean ± SD. Scale bars, 4 μm.

To test if SMAC could monitor circulating PSA in human blood samples, we measured PSA levels in plasma samples from prostate cancer patients (Table S1) as well as from control healthy male and female blood donors (Fig. 2G). We found that in most cases SMAC detected circulating PSA in prostate cancer patients at abnormally high levels (~10-100 pM) compared to baseline PSA levels in healthy male donors (~100 fM) (Fig. 2G). In contrast, conventional ELISA required >10-fold greater plasma volume to detect circulating PSA from prostate cancer patients, and could not detect basal PSA levels in control male blood donors (Fig. S5), which is consistent with prior studies (*4*). We also found that one prostate cancer patient had abnormally low circulating PSA levels (Fig. 2G and Fig. S5); it has been observed that ~10% of prostate cancer patients have very low circulating PSA levels (*19–21*), which correlates with poor prognosis (*22*). These results illustrate the clinical utility of SMAC for established, secreted biomarkers and demonstrate the first single-molecule imaging-based blood test, paving the way to applications of single-molecule imaging in non-invasive profiling of disease-associated proteins.

We next used SMAC to characterize membrane-bound proteins shed into the blood. Programmed death-ligand 1 (PD-L1) (*23*) is a membrane-bound immune checkpoint mediator which inhibits immune responses in a variety of disorders spanning from cancer to infection (*24–26*) and has recently been found in human blood (*27, 28*). PD-L1 antagonists have shown promise in treating chronic virus infection (*29*) as well as multiple cancer types (*30, 31*). A non-invasive approach to predict likelihood of benefit from immune checkpoint blockade could help tailor clinical management to individual patients (*32*). We thus employed SMAC to determine the level of circulating PD-L1 in human blood (Fig. 2H).

We first developed SMAC to detect purified PD-L1 down to aM concentrations (LOD of 607 aM) and with a six-log dynamic range (Fig. 2I, J and Fig. S6A). By comparison, the detection limit of conventional ensemble methods such as ELISA was 10^4^-fold higher (~10 pM) and had a two-log dynamic range (Fig. S6B). We next applied SMAC to characterize circulating PD-L1 molecules in patients with a chronic virus infection-induced disease: human papillomavirus (HPV)-associated cervical high-grade squamous intraepithelial lesions (*33*) (HSIL; Table S2). Baseline circulating PD-L1 levels spanned from 50-300 fM in control blood donors (Fig. 2K). By contrast, five of six HSIL patients had blood PD-L1 levels >300 fM, and two of these patients had levels >500 fM (Fig. 2K). By comparison, ELISA could detect circulating PD-L1 in only one of the six HSIL patients (Fig. S7). Circulating PD-L1 was likely increased in a subset of HSIL patients because HPV infection and T cell-mediated local inflammation together induce tissue PD-L1 gene expression (*34*). The heterogeneity in PD-L1 levels is consistent with previously reported percentages of ectopic PD-L1 expression in tissue from HPV-associated lesions (*35, 36*). These results introduce opportunities for single-molecule imaging to investigate the role of circulating immune checkpoint mediators as well as other membrane-tethered proteins in human diseases.

Having demonstrated the detection of extracellular proteins, including secreted and membrane-bound, proteins in blood using SMAC, we next turned to the detection of rare intracellular proteins shed from disease sites into the blood. Current blood tests target extracellular proteins as they are easily accessible (*37*), but many of these proteins are also found in the blood of healthy people (*38*). In contrast, certain intracellular proteins, in particular those that promote oncogenic transformation—such as mutant or viral oncoproteins, as well as mutant tumor suppressor proteins—are exclusively expressed by diseased cells and hence would be more accurate biomarkers than extracellular proteins. While we and others have observed the release of intracellular proteins from cultured cancer cells (*39*) (Fig. S8A), their presence in blood has not been reported, likely because they lie amidst a plethora of other circulating proteins and are too rare to be quantified by current methods. The idea of identifying circulating intracellular proteins, such as mutant proteins, has remained an elusive goal (*40*).

We first assessed if intracellular proteins from tumor cells are shed into the bloodstream using an animal model in which tumor cells (TC-1 (*41*)) are engineered to express cytoplasmic GFP (cytoGFP) as a prototype intracellular protein (Fig. 3A). We inoculated mice with cytoGFP^+^ tumor cells in either subcutaneous or mucosal (buccal) tissue and were able to detect cytoGFP in the blood of these mice within one week after tumor challenge (Fig. 3B). The concentration of circulating cytoGFP ranged from 1 fM to 1 pM (Fig. 3B), which was in most cases below the ELISA detection limit (Fig. S3B). We then induced mice with spontaneous tumor by electroporating into buccal tissue DNA vectors encoding oncogenes (*42*) (*Ras^G12V^* and *p53* shRNA) and the intracellular biomarkers cytoGFP (*cyto-gfp*) and luciferase (Fig. S8B). We were able to monitor the accumulation of cytoGFP in serum from these mice (from ~1 fM to ~1 nM), which paralleled tumor burden as measured by buccal luminescence imaging of luciferase activity (Fig. 3C and Fig. S8C). Serum *cyto-gfp* DNA was not detectable by quantitative PCR (qPCR) even when the tumor reached >5 mm diameter (Fig. S8D, E), likely due to the low copy number and labile nature of DNA in the blood (Fig. S8F). These results are consistent with the typically low levels of circulating tumor DNA (ctDNA), especially in early-stage cancer (*43–46*). Using SMAC to follow the release of cytoGFP protein over time, we found that circulating cytoGFP levels closely mimicked luminescence imaging of tumor onset and progression (*47*) beyond week two, once the initial circulating cytoGFP peak (due to electroporation-mediated tissue damage) had waned (Fig. 3D and Fig. S9). These results were confirmed by cross-correlation analysis (Fig. 3E). Of note, serum cytoGFP was imperceptible by ELISA even 42 days after tumor was induced (Fig. S10). These results underscore the potential of single-molecule imaging to study fundamental disease processes in animal models, as well as to identify rare intracellular proteins in the blood for early disease detection.

**Fig. 3.**
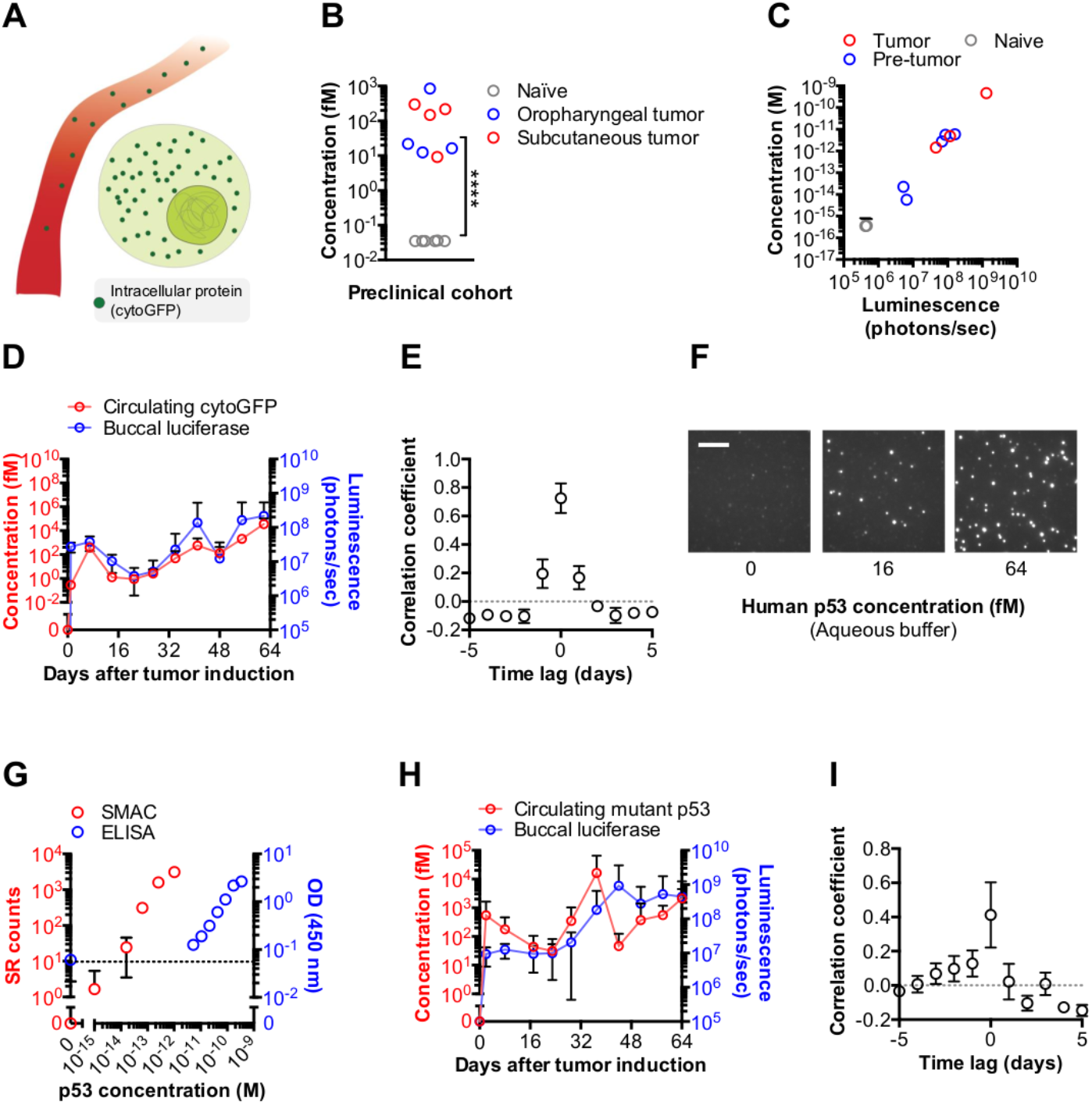
Detection of cytoplasmic and nuclear proteins in blood by single-molecule imaging. (A) Schematic diagram of intracellular cytoGFP release from a tumor cell (lime) into a blood vessel (red). (B) SMAC quantification of serum cytoGFP levels in naïve mice (gray circles; *n* = 8) as well as tumor-bearing mice one week after oropharyngeal (blue circles; *n* = 4) or subcutaneous (red circles; *n* = 4) injection of cytoGFP^+^ tumor cells (TC-1). (C-E) To induce a spontaneous cytoGFP^+^ tumor, mice (*n* = 10) were administered with DNA encoding Ras^G12V^, p53 shRNA, cytoGFP, and luciferase. Graph depicting the relationship between tumor luciferase and serum cytoGFP concentrations assessed by SMAC at an endpoint of more than two months (C) or throughout the first two months (D). In (C), tumor-induced mice that displayed a grossly visible tumor were labeled ‘tumor’ (red circles), while those that did not were labeled ‘pre-tumor’ (blue circles). Using the kinetics data in (D), the time correspondence between serum cytoGFP levels and tumor burden was determined by cross-correlation analysis (E). (F) SMAC images of purified human p53 at fM concentrations in aqueous buffer. (G) Comparison of the sensitivity of SMAC with shape analysis (SR counts, red circles) and ELISA (OD_450 nm_, blue circles) using purified human p53. The dotted line indicates the ELISA detection limit. (H, I) To stimulate a spontaneous tumor carrying mutant human p53, mice (*n* = 10) were administered with DNA encoding human p53^R175H^, Ras^G12V^, and luciferase. Time course (H) and cross-correlation (I) plots depicting the relationship between tumor luciferase and serum mutant p53 levels measured by SMAC. For cross-correlation plots, each unit time lag is around five days. All data are expressed as mean ± SD. \*\*\*\**P*<0.0001. *P*-values are from a two-sided unpaired *t*-test. Scale bar, 4 μm.

To investigate the release of rare intracellular proteins in a clinically important system, we focused on the transcription factor p53 since it is a well-established tumor suppressor and the most commonly altered protein in human cancers (*48*). We developed SMAC to detect fM levels purified human p53 protein in an aqueous buffer (LOD of 12 fM), ~10^4^-fold below the ELISA limit (Fig. 3F, G). In cancer cell lines carrying different mutant p53 variants, we detected substantial levels of p53; by contrast, we detected essentially no p53 in cell lines with wildtype (wt) p53 (Fig. S11A). These results reflect the enhanced stability of mutant p53 relative to wt p53, as the latter undergoes rapid degradation by proteasomes (*49, 50*). Note that we used anti-p53 antibodies that theoretically recognize total p53, including wildtype and mutant variants. However, because only the mutant form of p53 is detectable in cell lines, in subsequent experiments we interpreted the presence of p53 in samples as ‘mutant pattern’ p53. We were able to observe mutant p53 release into the extracellular milieu from as few as 300 human ovarian cancer cells (OVCAR3) cultured overnight (Fig. S11B).

We optimized SMAC with shape analysis to correct background signals for p53 spiked into serum (Fig. S11C). To characterize mutant p53 proteins shed into the bloodstream in an animal model, we induced tumor formation in mice by co-delivery of DNA encoding human mutant p53^R175H^, Ras^G12V^, and luciferase into the buccal mucosa using electroporation. We measured serum mutant p53 proteins in these mice over time by SMAC with shape analysis. Circulating mutant p53 levels rose in parallel with tumor progression (from ~75 fM two weeks after tumor onset to ~2 pM after two months), even in the case of tumor metastasis, as assessed by luminescence imaging (Fig. 3H, I and Fig. S12A, B).

We next utilized SMAC to identify mutant p53 proteins in the blood of patients with high-grade ovarian cancer (HGOC; Table S3) (Fig. 4A), since the tumor from >96% of HGOC patients contains mutations in the *TP53* gene (*51*). Using normal human plasma spiked with purified p53, we verified that SMAC with shape analysis maintained fM baseline sensitivity (LOD of 35 fM) for p53 in human blood yet corrected >96% of background errors (Fig. 4B, C and Fig. S13). We detected mutant pattern p53 molecules in ~60% of plasma samples from HGOC patients with disseminated (FIGO stage III) disease (ranging from <10 fM to 1 pM) but in none of the samples from age-matched control female blood donors (Fig. 4D). We reasoned that for patients with undetectable circulating p53, host autoantibodies may have depleted p53 proteins (*52*) or blocked their capture (Fig. 4A). We therefore developed SMAC to identify autoantibodies against p53. SMAC exhibited >10^3^-fold greater sensitivity than existing assays and could detect autoantibodies in picoliter (10^−12^ l) volumes of human blood (Fig. 4E, F). Using SMAC, we measured abundant amounts of plasma anti-p53 autoantibodies in 43% of the cohort of HGOC patients (~10^3^-fold in excess of circulating mutant p53 levels) but not from any healthy donors (Fig. 4G). Interestingly, the presence of circulating mutant p53 and its autoantibodies appeared anti-correlated (Fig. 4H), suggesting that host immune responses might have cleared mutant tumor antigens from the blood or that autoantibodies disrupted capture of p53 (*53, 54*). Altogether, SMAC detected circulating p53 protein or abundant anti-p53 autoantibodies in 86% of HGOC patients and in no healthy subjects (Fig. 4H).

**Fig. 4.**
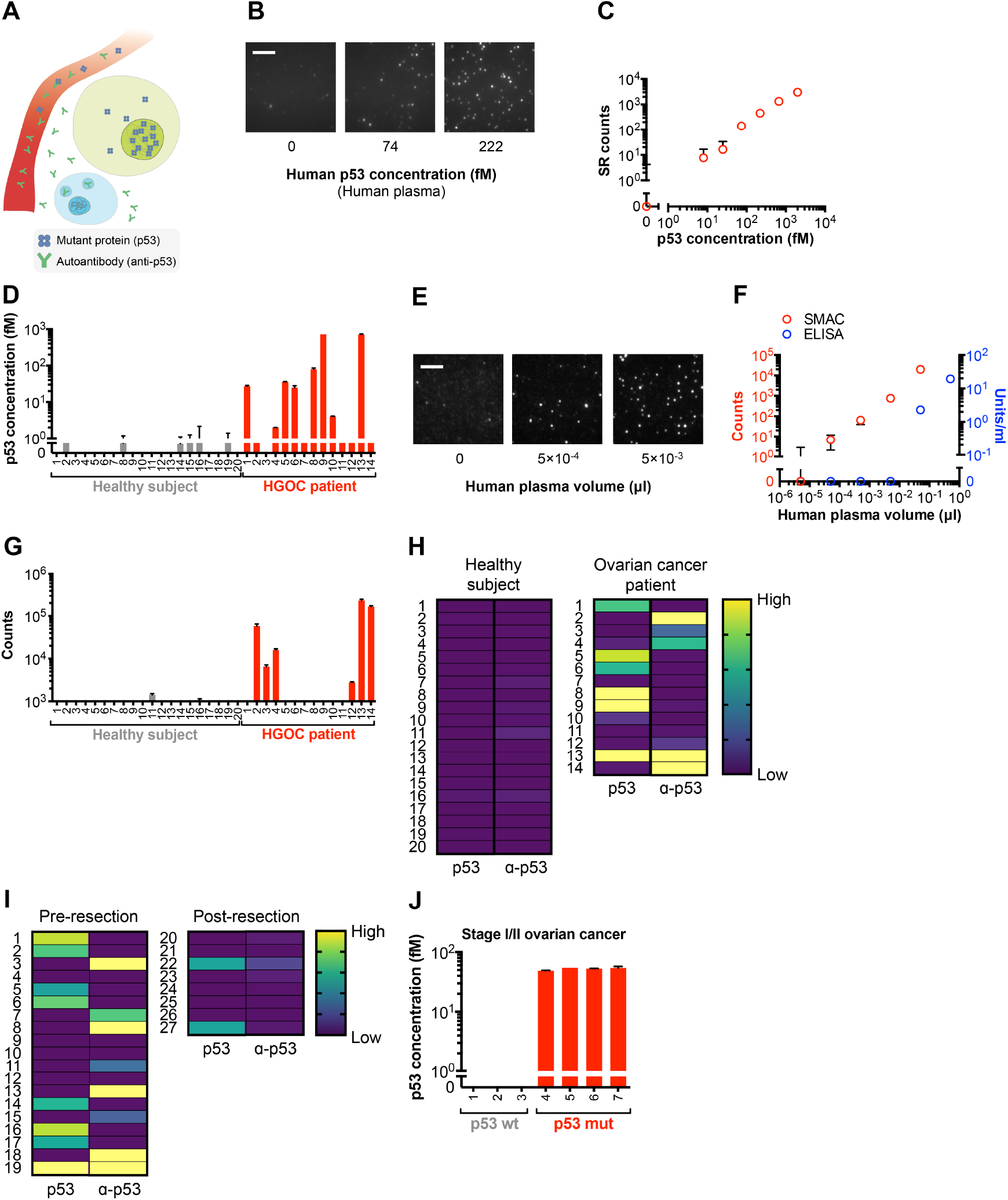
Detection of circulating mutant proteins and autoantibodies in blood by single-molecule imaging. (A) Schematic diagram depicting release of nuclear p53 from a tumor cell (lime) and anti-p53 autoantibodies from a tumor-specific B cell (aqua) into a blood vessel (red). SMAC images (B) and shape analysis (C) of purified human p53 added at fM concentrations in human plasma. (D) Shape analysis of circulating mutant p53 levels in plasma from high-grade ovarian cancer (HGOC) patients and healthy female blood donors. (E) SMAC images of endogenous anti-p53 autoantibodies in different plasma volumes, from the microliter (μl; 10^−6^ l) to picoliter (pl; 10^−12^ l) range, in an HGOC patient. (F) Comparison of the sensitivity of SMAC (counts, red circles) and ELISA (units/ml, blue circles) using human anti-p53 autoantibodies in human plasma. (G) Quantification of endogenous plasma anti-p53 autoantibodies from HGOC patients and healthy female blood donors; same cohort as in (D). (H) Heat map depicting the relative levels of circulating mutant p53 or anti-p53 autoantibodies in the blood of HGOC patients and healthy blood donors; same cohort as in (D) and (G). (I) Heat map depicting the relative levels of circulating mutant p53 or anti-p53 autoantibodies in an independent cohort of FIGO stage III ovarian cancer patients with p53-mutant tumors either before or after surgical resection. (J) SMAC analysis of circulating mutant p53 levels in early-stage (FIGO stage I/II) ovarian cancer patients with either p53-wildtype or p53-mutant tumors. Data for individual human plasma samples (D, G, J) are expressed as mean ± SE; all other data are expressed as mean ± SD. Scale bars, 4 μm.

We next characterized the levels of circulating mutant p53 and anti-p53 antibodies in an independent validation cohort of ovarian cancer patients with well-defined clinical information and p53 mutation status presenting at various pathologic stages, including early-stage (FIGO stage I and II) disease, either pre- or post-surgical resection. These samples were from the same cohort of patients included in the recent study describing the CancerSEEK technique by Vogelstein and coworkers (*55*). Of note, detection of ovarian cancer at an early stage, when surgical resection may be curative, remains a critical challenge in the field, as less than 20% of ovarian cancer cases are identified at stage I or II (*56*). Among ovarian cancer patients with stage III disease—all of whom carried mutant p53 within the tumor (Table S4)—79% (15 of 19) had either circulating mutant p53 (8 of 19) or anti-p53 autoantibodies (8 of 19) prior to surgical resection (Fig. 4I and Fig. S14). Only one patient had both mutant p53 and anti-p53 autoantibodies within the blood. Together, these results suggest that the presence of circulating mutant p53 and its autoantibodies is anti-correlated. Two of the four patients without circulating mutant p53 or anti-p53 autoantibodies had non-serous ovarian cancer (carcinosarcoma and undifferentiated carcinoma) (Table S4). The role of p53 alterations as a driver of serous ovarian cancer has been well-established (*51*), but its role in other histologic types of ovarian cancer remains unclear. Among stage III ovarian cancer patients previously treated by surgical resection, only 25% (2 of 8) had mutant p53 in the plasma (Fig. 4I and Fig. S14), suggesting that circulating mutant p53 levels serve as an index of tumor progression following therapy.

Included in the validation cohort were seven ovarian cancer patients with early-stage (stage I and II) disease (Table S5). We identified circulating mutant p53 protein in four of these patients (Fig. 4J), all of whom also carried corresponding genetic alterations in *TP53* within their tumors (Table S4). However, the three patients without circulating mutant p53 protein had tumors that lacked *TP53* alterations (Table S4). Interestingly, all early-stage ovarian cancer patients with p53-mutant tumors displayed circulating mutant p53 but not anti-p53 autoantibodies, suggesting that autoantibodies have not yet formed against p53 in patients with early-stage disease. Together, these data indicate that intracellular mutant driver proteins, such as p53, are shed into the bloodstream early on in tumorigenesis, and the analysis of these proteins by single-molecule imaging, in conjunction with tissue-specific biomarkers, may facilitate earlier, more accurate detection and diagnosis of disease, when surgical resection would have more clinical benefit.

The potential applications of ultra-sensitive single-molecule protein imaging extend beyond quantification of target proteins. In fact, SMAC can be used to investigate biochemical properties (e.g., secondary modifications, structural changes, aggregation status) of individual proteins-of-interest within a population, as well as unique combinations of proteins in macromolecular complexes. Because disease-associated proteins often differ between patients and healthy people not only in their total amount but also in their biochemical features, SMAC adds a new dimension to the information obtainable from existing methods.

To explore this avenue, we developed SMAC to investigate the aggregation status of p53 complexes in test samples. Of note, certain conformational mutants of p53 have been shown to self-assemble into high-order complexes within tumor cells, and these mutants have been correlated with aggressive disease (*57*). Thus, the ability to identify these conformational p53 mutants could improve disease detection and management. To test the idea that SMAC could distinguish between conformational p53 mutants, we generated p53 mutants which have been reported to self-assemble into large complexes (p53^R175H^) or remain as monomers (p53^L344P^) (Fig. S15A). We fused these mutants to the GFP reporter protein. We then added the recombinant mutant or wildtype p53 into buffer at different concentrations and examined them by SMAC. We found that the p53 conformational mutants produced different combinations of the number and intensities of fluorescent spots despite equal protein amounts (Fig. S15B, C). For example, at the same total p53 concentration, p53^L344P^ had a large number of low intensity spots; by contrast, p53^R175H^ had fewer spots, but these spots were of high intensity (Fig. S15B, C).

We characterized the intensity distributions of the fluorescent spots and measured the percentage of aggregates (defined as spots greater than or equal to tetramer) in each group of conformational mutants (Fig. S15D-F). At the same p53 concentration, p53^R175H^ had the largest percentage of aggregates, followed by the wildtype group and then p53^L344P^ (Fig. S15E). Also, the relationship between the percentage of aggregates and fluorescent spot number was different among each group of conformational mutants (Fig. S15F). We found that the aggregation-prone p53^R175H^ mutant had the widest intensity distribution of fluorescent spots, followed by wildtype p53, and then the monomeric p53^L344P^ mutant for a given spot number (Fig. S15D). To quantify these data, we calculated the Fano factor of fluorescent p53 spots (defined as the variance in intensity divided by the mean intensity; see Methods for details) as a relative index of the aggregation status (and therefore the likely mutation status). The p53^R175H^ mutant had the greatest change in Fano factor per unit change in spot number, followed by wildtype p53 and then the p53^L344P^ mutant (Fig. S15G). These data indicate that SMAC can reveal conformational properties of disease-associated proteins and open up the possibility of using single-molecule imaging to investigate the structural properties of mutant p53, as well as other disease-associated proteins, in clinical samples.

In summary, our results illustrate broad applications of single-molecule imaging to characterize disease-associated secreted, membrane, and intracellular proteins in the blood, opening new avenues to detect, diagnose, and study disease. We are in the process of converting the SMAC technology into a platform that can be broadly applied in clinical practice. To achieve this, we are developing an integrated device that combines the microfluidic handling, single-molecule imaging, and data analysis components of SMAC. The total run-time of the SMAC assay is approximately 4 hours, including target protein capture, detection antibody incubation, and single-molecule imaging, which is in-line with the time required for existing protein detection methods. Furthermore, while we have found that the sensitivity of SMAC correlates with affinity of the capture/detection antibodies employed, detection limits in the fM and sub-fM range are attained with antibodies with dissociation constants (KD) ~10 nM or lower, which can be achieved for the vast majority of target proteins with modern antibody production technologies. Thus, we believe the SMAC platform can be readily adapted for disease detection, diagnosis, and monitoring in the clinical setting.

While it introduces single-molecule imaging into the clinical arena and serves as a powerful tool for disease profiling, there are limitations of the SMAC system in its present form. First, the TIRF microscope imaging is performed sequentially, with a total acquisition time of approximately 5 minutes per sample. Thus, the scale at which samples can be run is currently limited. Second, for target proteins that are disease-specific and absent in control samples (such as mutant tumor proteins), control plasma from healthy blood donors can be used in the data analysis algorithm to normalize for background fluorescence in the blood (such as from substances that cause autofluorescence or non-specific detection antibody binding). However, for target proteins that are also present to varying degree in control samples, such as tumor-associated proteins, buffer solution (rather than plasma) must be used as a negative control. For these types of target proteins, the analysis algorithm unable to separate true signal from background in plasma.

Altogether, the insight gained from SMAC may shed light on pathologic processes, such as dysfunctional signaling pathways, gene expression networks, or immune responses unfolding within disease and point to effective therapies. The platform described here may be adapted to investigate unique biochemical, conformational, and structural features of proteins-of-interest in the blood. The design of SMAC can also be readily converted into multiplex and high-throughput formats to enable large-scale, single-molecule profiling of proteins in human disease.

## Acknowledgments

We are grateful to Dr. Christopher Bohrer and Mr. Huy Vo for assistance with microfluidic device design and fabrication. We thank Dr. TC Wu (T.C.W.) and Dr. Bert Vogelstein for their insightful input on the manuscript. We also thank Dr. Xinxing Yang, Dr. Ryan McQuillen, and Dr. Kelsey Bettridge for helpful discussion;

## Author contributions

C.P.M., S.C.W., and C.F.H. designed research; C.P.M., S.C.W., and Y.P.S. conducted single-molecule imaging experiments and data processing. Y.P.S. and S.H.T. performed mouse experiments and ELISA. L.H., Y.C.T., A.A.W. conducted protein purification; R.B.S.R. supplied human plasma samples; C.P.M., S.C.W., R.B.S.R., J.X. and C.F.H. wrote the paper;

## Data and materials availability

Key data to support the conclusions of this study are available in the main figures and Supplementary Materials. Additional substantiating data, including raw data for all figures, are available upon request. Programming codes for the SMAC analysis algorithms are available upon request. Correspondence and requests for materials should be addressed to.

## Supplementary Materials

Materials and Methods

Figures S1-S15

Tables S1-S5

References (*58, 59*)

