## Supplementary_materials for "Protein detection in blood with single-molecule imaging"

**This PDF file includes:**

Materials and Methods
Supplementary Text
Figs. S1 to S15
Tables S1 to S5

### Materials and Methods

Materials

Polydimethylsiloxane (PDMS) elastomer for synthesis of the single-molecule microfluidic capture device was purchased from Dow Corning. 22×22 mm borosilicate cover glass (Thermo Fisher Scientific) served as the substrate for the capture surface. Surface passivation required the following reagents: *N*-(2-aminoethyl)-3-aminopropyltrimethoxysilane (United Chemical Technologies), Alconox (Alconox, Inc.), methanol (Fisher Scientific), acetic acid (Sigma-Aldrich), sodium bicarbonate (Sigma-Aldrich), biotin-mPEG-succinimidyl valerate MW 5,000 (biotin-mPEG-SVA) (Laysan Bio), and mPEG-succinimidyl valerate MW 5,000 (mPEG-SVA) (Laysan Bio). For GFP detection, biotinylated anti-GFP antibodies (clone RQ2, MBL) were used. For PSA detection, biotinylated (BAF1344, R&D Systems) and Alexa Fluor 488-conjugated (clone 8301, Medix Biochemica) anti-human PSA antibodies were used. For PD-L1 detection, biotinylated (BAF156, R&D Systems) and Alexa Fluor 555-conjugated (clone 28-8, Abcam) anti-human PD-L1 antibodies were used. For human p53 detection, biotinylated (BAF1355, R&D Systems) and Alexa Fluor 555-conjugated (clone E47, Abcam) anti-human p53 antibodies were used. For human p53 detection in mice, biotinylated (BAF1355) and Alexa Fluor 488-conjugated (FL-393, Santa Cruz Biotechnology) antibodies were used. For detection of human anti-p53 autoantibodies, Alexa Fluor 555-conjugated anti-human IgG (H+L) cross-absorbed secondary antibodies (A-21433; Thermo Fisher Scientific) were used. Purified recombinant GFP (Cell Biolabs), human PSA (R&D Systems), human PD-L1 (R&D Systems), and human p53 (R&D Systems) were used to generate standard curves. BSA (New England BioLabs) and polyoxytheylene (20) sorbitan monolaurate (Tween-20; Thermo Fisher Scientific) were used for sample wash and dilution in single-molecule experiments. ELISA for GFP, PSA, PD-L1, p53, and anti-p53 autoantibodies was performed with GFP ELISA Kit (Cell Biolabs), PSA Quantikine ELISA Kit (R&D), PD-L1 Quantikine ELISA Kit (R&D), p53 SimpleStep ELISA Kit (Abcam), and Mesacup Anti-p53 Test (MBL International), respectively, according to manufacturer instructions.

Antibody conjugation

Antibodies were labeled with biotin or organic fluorophores via NHS-reactive ester. For biotin conjugation, antibodies (0.1-1 mg/ml) were incubated with 50-fold molar excess of NHS-LC-biotin (Thermo Fisher Scientific) for 30-60 min and then isolated on 7 kD gel filtration columns (Thermo Fisher Scientific). For dye conjugation, antibodies were pre-captured on protein G magnetic beads (Thermo Fisher Scientific) and incubated with 10-fold molar excess of Alexa Fluor dye-NHS (Thermo Fisher Scientific) for 30-60 min. Free dye was washed out, and antibodies were further purified on 7 kD gel filtration columns. Degree of labeling and concentration of antibodies were measured by spectrophotometry.

Pairwise antibody screening

To select the best pair of capture and detection antibodies recognizing proteins-of-interest, candidate antibodies were each labeled with biotin or organic dye as described above. Pairwise combinations of these candidate antibodies were then evaluated by flow cytometry on microbeads. Biotinylated capture antibodies (1 μg) were incubated with streptavidin M-280 magnetic Dynabeads ($10^5$ beads per sample; Thermo Fisher Scientific) for 30 min. The beads were washed with PBS and incubated with or without purified target proteins for 30 min. Beads were washed with PBS and incubated with different dye-labeled detection antibodies (1 μg) for 30 min. These procedures were carried out in 50 μl total volume at 25°C with constant mixing. Beads were then washed, resuspended in PBS (500 μl), and interrogated by flow cytometry on a FACSCalibur device (BD Biosciences). The capture/detection efficiency for each pair of antibodies was calculated based on the shift in mean fluorescence intensity in the presence versus absence of target proteins. Antibodies that yielded the greatest shift were considered to have superior performance.

Purification of recombinant human p53

Human p53 was generated in-house for anti-p53 autoantibody detection experiments. Plasmid encoding human p53 (hp53; Addgene) was inserted into the pET28a bacterial expression vector. *hp53* was first amplified by PCR with the following primer set: 5' AAAGGATCCATGGAGGAGCCGCAGTCAGA 3' and 5' AAAGAATTCCAGGTGGCTGGAGTGAGCCC 3'. The PCR product was cloned into the BamHI/EcoRI sites of the pET28a vector to create pET28a-hp53. This plasmid was transformed into *E. coli* BL21 competent cells (Novagen). Protein expression was induced with 1 mM isopropyl-b-D-thiogalactopyranoside (IPTG) at 37°C for 5 hr. Bacteria were lysed, and the soluble fraction was collected. Recombinant protein was purified by affinity chromatography on Ni-NTA agarose (Qiagen) per manufacturer instructions. Purified p53 was verified by 10-15% gradient sodium dodecyl sulfate polyacrylamide gel electrophoresis (Bio-Rad) and Coomassie Brilliant Blue (Thermo Fisher Scientific) staining, dialyzed with PBS, and stored at -80°C in PBS containing 20% glycerol.

Cells

TC-1 cells were previously generated in our laboratory and have been reported[42]. For experiments involving cytoGFP, TC-1 cells were retrovirally transduced with a *cyto-gfp* DNA expression cassette. LnCaP human prostate cancer cells were obtained from ATCC. Human cells without p53 (BHK21) and with mutant p53 (CFPAC-1 (C242R), OVCAR3 (R248Q), and TOV-112D (R175H)), cells with wildtype p53 (MCF-7, MCF-10), and HEK 293T cells were from ATCC. Cells were cultured in RPMI-1640 medium or DMEM (Thermo Fisher Scientific) with 10% FBS in the absence of phenol red and maintained under 5% $CO_2$ atmosphere. Lysate was prepared using commercial lysis buffer (Abcam). Briefly, cells were harvested and resuspended in lysis buffer at a stock concentration of $10^4$ cells per μl. The resultant solution was centrifuged at 10,000×*g* at 4°C for 10 min. Target protein concentration in the stock lysate was determined by ELISA. For single-molecule imaging experiments, lysate was diluted $10^3$-$10^5$× in 'SMAC buffer' (10 mM Tris-HCl pH 8.0, 50 mM NaCl, 0.05% Tween-20) with 0.1 mg/ml BSA. For experiments involving supernatant, conditioned medium was collected from cells cultured for 12-24 hr and centrifuged at 1,000×*g* for 5 min. The resultant supernatant was passed through 0.22-μm filters to further remove debris. The number of viable cells was determined using an automated cytometer (Countess II, Invitrogen) with trypan blue dye exclusion.

ELISA

Enzyme-linked immunosorbent assay (ELISA) was performed according to the manufacturer's instructions. Briefly, plasma or cell lysate (50 μl) was added to sample diluent (50 μl). The mixtures were then added to antibody pre-coated plates and incubated at room temperature for 2 hours. The plates were washed with PBS containing 0.05% Tween-20, and HRP-conjugated antibodies were then added to the plate and incubated at room temperature for 2 hours. The plates were washed as above and then developed with TMB substrate solution. The reaction was terminated with stop solution containing 1 M phosphoric acid. The signals in the plates were measured at 450 nm wavelength using a microplate reader (Bio-Rad).

Mice

6- to 8-week old female C57BL/6 and immune-deficient athymic nude (*Foxn1*$^{-/-}$) mice were obtained from the National Cancer Institute. C57BL/6 mice were used for experiments in which tumor cells were directly inoculated. *Foxn1*$^{-/-}$ transgenic mice were used for experiments in which a spontaneous tumor was induced by oncogene delivery. All animal procedures complied with protocols approved by the Johns Hopkins Institutional Animal Care and Use Committee and with recommendations for the proper use and care of laboratory mice.

Transplanted tumor challenge

C57BL/6 mice were injected with cytoGFP-transduced TC-1 cells ($10^5$ cells per animal) in the flank (subcutaneous tissue) or buccal mucosa. At one week after tumor challenge, whole blood was collected from the tail vein and processed into serum for downstream experiments. Tumor growth was monitored by visual inspection, palpation, and digital caliper measurement.

Spontaneous tumor induction

*Foxn1*$^{-/-}$ transgenic mice were injected in the buccal mucosa with a plasmid DNA cocktail encoding (1) mutant Ras$^{G12V}$, (2) SB13 transposase, (3) firefly luciferase, and (4) either anti-p53 shRNA carrying a GFP expression cassette or mutant human p53$^{R175H}$ (10 μg of each plasmid diluted with PBS to 30 μl total volume); plasmids were acquired from Addgene. Immediately afterward, mice received electroporation (eight pulses of 72 V, 50 ms duration, and 200 ms interval) at the injection site with an ECM830 device (BTX). Tumor burden was monitored over time by whole body luminescence imaging of luciferase activity with an IVIS Spectrum device (PerkinElmer) following intraperitoneal D-luciferin (Promega) injection. At defined time points after tumor induction, whole blood was collected from the tail vein and processed into serum for downstream experiments.

Cross-correlation analysis

To characterize the time relationship between tumor progression and fluctuations in circulating target protein levels in mice, the covariance was calculated between time series of error-corrected single-molecule serum target counts and luminescence photon counts. The covariance coefficient was computed via the ‘xcov’ function in Matlab (MathWorks). The time lag was narrowed down to five units, with each lag unit corresponding to approximately five days.

Human subjects

Blood samples were obtained from volunteer patients previously diagnosed with prostate adenocarcinoma ($n$ = 5), high-grade cervical intraepithelial lesions ($n$ = 6), and ovarian cancer ($n$ = 48) who underwent clinical evaluation and management at the Johns Hopkins Hospital, Baltimore, MD, USA. Descriptions of the clinical characteristics of individual patients are provided in Tables 1-3. Human studies were approved by the Johns Hopkins University Institutional Review Board (protocol number: IRB00169055). Plasma from healthy human volunteers was acquired from Innovative Research, processed from whole blood in dipotassium ethylenediaminetetraacetic acid (K2-EDTA; BD Biosciences).

Human plasma preparation

Whole blood was drawn from test subjects and anticoagulated with K2-EDTA. Samples were processed within 4 hr after collection. Blood samples were diluted with an equal volume of 1× HBSS (Corning) and added slowly on top of Lymphoprep solution (15 ml; Stem Cell Technologies) in 50-ml conical tubes (Corning). Samples were centrifuged at 1,200×*g* for 10 min at room temperature. The top layer was harvested as plasma and stored at -80°C.

Single-molecule capture surface passivation

Borosilicate coverslips of 130-170 μm thickness and 22×22 mm area served as the substrate for the capture surface. Coverslips were first cleaned in 1% Alconox with sonication for 10 min, washed with Milli-Q water (Millipore) for 10 min, and dried with filtered air. Coverslips were exposed to high power atmospheric plasma using a PE25-JW device (Plasma Etch) for 5 min for surface cleaning and activation, and then immediately dipped in methanol containing 1% *N*-(2-aminoethyl)-3-aminopropyltrimethoxysilane and 5% glacial acetic acid. Coverslips were washed thoroughly with methanol and Milli-Q water, and then dried with filtered air. Coverslips were conjugated with biotin-mPEG-SVA (0.3 mg) in 10 mM sodium bicarbonate (pH 8.5) for 6 hr in a sandwich arrangement. The glass surface was then conjugated with a mixture of biotin-mPEG-SVA (0.3 mg) and PEG-mSVA (16 mg, 1:50 mass ratio) for 12 hr in a sandwich arrangment. After passivation, coverslips were washed with Milli-Q water and dried as described above. Coverslips were transferred to a clean container, vacuumed, flushed with pure nitrogen, sealed with paraffin film, and stored at -20°C. Tween-20 was added into SMAC buffers during downstream experiments to further block the surface.

Microfabrication of the SMAC chip enclosure

A master template for the device enclosure was synthesized by photolithography. Briefly, a silicon wafer was rinsed with acetone and isopropanol and then dehydrated at 200°C for 15 min. The wafer was exposed to high power oxygen plasma (100 W for 3 min at 300-500 mTorr oxygen pressure) using a PE II-A apparatus (Technics) to promote photoresist adhesion. SU-8 photoresist 2050 (MicroChem) was spin-coated onto the wafer to 100 μm thickness. The wafer was then soft-baked (65°C/95°C) for 5 min and exposed to UV light in an EVG620 mask aligner (EVG) loaded with a mask printed at 32,512 DPI resolution (Fineline Imaging). The wafer was then hard-baked (65°C/95°C) for 15 min. The first layer of the microfluidic device consisted of the main channel with side boxes while the second layer contained arrays of staggered herringbone grooves. After all layers of photoresist were deposited, the wafer was developed under ultrasonic agitation to yield a master template for synthesis of the silicone elastomer enclosure. To produce this enclosure, PDMS elastomer was mixed with curing agent in a 10:1 ratio (by weight), poured onto the patterned wafer, degassed, and incubated at 80°C overnight. The PDMS was then removed from the master, cut into individual devices, and bored with inlet/outlet tubing holes (750 μm diameter). The devices were washed in an ultrasonic bath with isopropanol for 20 min and then with Milli-Q water for 5 min. Devices were dried with filtered air.

Assembly of the SMAC chip

Prior to assembly, the uncoated side of the borosilicate coverslip was taped to an alignment guide imprinted with a two-dimensional replica of the flow channel. An elastomer cover microfabricated with μm-precision by photolithography to match the exact size and shape of the flow channel was then placed on the coated side of the coverslip at the position of the channel replica on the alignment guide. This cover protects the PEG/biotin-PEG layer from oxygen plasma bombardment during the assembly procedure. The coated coverslip surface with elastomer cover and PDMS enclosure were placed inside a PE-25JW plasma etcher and treated with oxygen plasma for 30 sec at 40 W RF power under 100 mTorr oxygen atmosphere. The elastomer cover was removed, and PDMS devices were then sealed to the coated side of the coverslip under a stereomicroscope with the alignment guide as a reference for the channel position. The microfluidic chip was incubated at 80°C for 3 min to drive the bonding to completion.

Preparation of the SMAC chip

Reagent introduction, removal, and wash steps were performed in parallel under automated flow actuated by a multi-channel peristaltic pump (Ismatec). The SMAC chip was connected to inlet and outlet non-shrinkable Teflon tubing (internal diameter 0.015 in; Weico Wire and Cable) and infused with 'SMAC buffer': 10 mM Tris-HCl pH 8.0, 50 mM NaCl, and 0.05% Tween-20. The inclusion of Tween-20 in the SMAC buffer further blocked non-specific protein absorption to the PDMS chamber and the capture surface. To evacuate any air trapped inside the PDMS channel, the chip was immediately degassed under vacuum for 1 min. The chip was equilibrated with SMAC buffer at 50 μl/min flow rate for 10 min. The chip was then incubated with NeutrAvidin (20 μl; 0.1 mg/ml; Thermo Fisher Scientific) in SMAC buffer for 10 min. The chip was washed with SMAC buffer (1 ml) at 500 μl/min and incubated for 30 min with biotinylated antibodies (2 μl; 0.1-1 mg/ml) in SMAC buffer with 0.1 mg/ml BSA ($SMAC^{BSA}$ buffer). The chip was then washed with $SMAC^{BSA}$ buffer (1 ml) at 500 μl/min and ready for sample circulation.

Sample circulation in the SMAC chip

Continuous oscillating flow was actuated by a multi-channel bidirectional AL-8000 syringe pump (World Precision Instruments) connected to the SMAC chip via a 26-gauge 1-cc syringe (BD Biosciences). The chip was connected at the other tubing port to the sample prepared in $SMAC^{BSA}$ buffer. For oscillating flow, the blood sample was diluted to anywhere between 2% to 50% with $SMAC^{BSA}$ buffer in 200-500 μl final volume. Note that although we used 200-500 μl final sample volumes in this study, the SMAC system can actually accommodate volumes up to 10 ml without significant loss in sensitivity because of its oscillating flow scheme and efficient target capture. By contrast, most other methods are unable to reliably detect proteins in sample volumes much greater than 100 μl. The syringe pump was programmed to carry out repeated infusion/withdrawal cycles at 500 μl/min for 2-4 hr. Afterward, the chip was washed with $SMAC^{BSA}$ buffer (1 ml) at 500 μl/min, incubated with fluorophore-labeled detection antibodies (1-10 nM) for 30 min, and then washed again with SMAC buffer (1 ml) at 500 μl/min. For circulation of clinical plasma samples, the prostate cancer patient samples were diluted 50 times. HSIL patient samples were diluted two times. HGSOC patient samples were diluted two times for p53 detection and $10^3$-$10^6$ times for anti-p53 autoantibody detection. For circulation of mouse samples, serum was diluted four times. All dilutions were carried out in $SMAC^{BSA}$ buffer. For experiments involving human samples, ~$10^4$-fold excess IgG matched to the isotypes of the capture and detection antibodies was further added to reduce non-specific binding.

Single-molecule TIRF microscopy

An objective-based TIRF setup was employed with a PlanApo 60× oil objective (Olympus) of high numerical aperture (1.45). While acquiring data, we also used a 1.6× field lens to capture single-molecule images at 96× total magnification. The incident laser angle was adjusted to full TIRF mode with a prism. Flow channels in the SMAC chip were identified under brightfield illumination. An imaging region of 15×15 μm$^2$ was then set. An electron multiplying charge-coupled device camera (Andor) was programmed to capture a consecutive time stream of 500 frames with 50 ms exposure time under continuous laser excitation of 40-140 W/cm$^2$. Immediately prior to imaging, we measured laser power and TIRF angles to confirm that they were consistent. After imaging each region, the stage was displaced 80 μm down the length of the channel, and imaging was performed again as above. This process was repeated until at least 10 view fields were recorded per sample. Data were acquired with custom journals written in MetaMorph software (Molecular Devices).

Integrated intensity analysis. This method was used to quantify relatively abundant proteins-of-interest (e.g. >10 fM) across a wide dynamic range (6-9 logs). Single-molecule TIRF data were first recorded using a camera EM gain setting of 300. The EM gain of the camera (i.e., 300, 10, or 1) was chosen according to predefined criteria based on the standard curve for that protein. These criteria were set such that the florescence for the sample would fall within the linear range of the standard curve for that EM gain setting.

Single-molecule shape analysis

This method was used for rare proteins-of-interest to correct detection errors due to diffusive background and non-specific binding. The first 50-100 frames of each TIRF image were initially averaged to reduce background fluorescence from compounds arbitrarily deposited on the microfluidic capture chip, as these compounds bind weakly and rapidly dissociate from the chip. We measured the total number of fluorescent spots over 10 view fields in order to maximize sensitivity for rare target proteins. Single-molecule data were interpreted with the ThunderSTORM plug-in in ImageJ software (*58*). In ThunderSTORM, a wavelet filter was applied to remove noise and automatically identify fluorescent spots at a constant low threshold for each sample independent of the target protein. A low threshold setting was chosen in order to ensure that all potential target spots were selected regardless of variations in laser illumination intensity.

The major principle behind shape analysis is that, because target proteins are pulled down as complexes by multivalent antibodies via NeutrAvidin adapters, the shape of fluorescent spots, each represented in coordinates of ($I$; measured in number of photons) and diffraction-limited spot size (measured by the $\sigma$ of the Gaussian fitting of the spot), was non-identical between real signals and false signals from diffusive and non-specifically absorbed antibody molecules. Therefore, a raw SMAC image can be deconvoluted into its real and false (i.e. background) components. To do so, we converted each selected spot in the raw SMAC image into an $I$-$\sigma$ coordinate and sorted each spot into its respective bin in a 2D $I$-$\sigma$ histogram. Note that each bin $i$ of this histogram contains both real and false spots, adding up to a total of $T_i$ spots. The next step in the analysis is to determine the number of false spots in each bin. For each target protein under different conditions, we performed SMAC on control samples lacking the target protein. For experiments involving aqueous buffer and cell supernatant, SMAC$^{BSA}$ buffer and culture medium, respectively, were used as reference samples. For animal experiments, serum from naïve mice was used as reference samples. For experiments involving human blood, plasma from multiple independent healthy blood donors was used as reference samples.

The identified spots in these control images were also sorted into bins in a 2D $I$-$\sigma$ histogram. Note that each bin of this reference histogram contains only false spots. By running this assay and analysis procedure on a large set of reference samples (e.g. buffer only or blood samples from many individual healthy donors), the mean ($R_i^{\mu}$) and standard deviation ($R_i^{\mathrm{SD}}$) of the number of spots in each bin was calculated. To correct detection errors and compute the number of real spots ($C_i^{\mu}$) for each bin, the following formula was used: $C_i^{\mu} = T_i - \left(R_i^{\mu} + n \times R_i^{\mathrm{SD}}\right)$, where $n$ can be adjusted the control the maximum number of projected false spots in each bin. For this study, $n = 2$ was chosen, as statistically there is a <3% chance that the number of false spots in each bin exceeds $R_i^{\mu} + 2R_i^{\mathrm{SD}}$. The total number of real spots was reported as 'SR counts', which was calculated by summing $C_i^{\mu}$ for every bin of the test sample 2D histogram. The limit of detection (LOD) was calculated using the following formula (*59*): LOD = SR $\text{counts}_{\text{control}} + 1.645 \times \text{SD}_{\text{control}} + 1.645 \times \text{SD}_{\text{min}}$, where $\text{SD}_{\text{min}}$ is the lowest protein concentration in which the mean number of SR counts exceeds its standard deviation. Note that with shape analysis, the probability that a sample would have SR counts ≥1 by chance alone is <3%. To further reduce the false positive rate for circulating mutant protein detection in clinical samples, we only considered samples to be positive at SR counts >3.

Analysis of protein aggregation

Various GFP-fused mutant p53 ($\text{p53}^{\text{R175}}$, $\text{p53}^{\text{L344P}}$) or wildtype p53 were expressed in a p53-deficient cell line, BHK21. The concentrations of p53 in the cell lysates were normalized by on denaturing SDS-PAGE. Serial dilutions were performed based on these normalized concentrations. Individual fluorescent spots were initially selected on the first imaging frame using ThunderSTORM for the various p53 protein conformational variants at different concentrations. Because the conformational variants produce spots with distinct intensity distributions, intensity histograms were generated from these spots, and fixed Gaussian fitting was applied to identify curves for different structural populations of p53 (e.g., monomer, dimer, tetramer, octamer). The areas under the curve for populations greater than or equal to tetramer were integrated and defined as aggregates. The structural compositions of p53 in each sample were hence determined based on the relationship between percentage of aggregates and spot number. Note that although spot number is directly influenced by protein concentration, the relationship between protein concentration and spot number varies depending on p53 conformation (for example, monomers yield a larger number of imaging spots than aggregates for any given total p53 protein concentration). The spread of intensity distributions allowed us to distinguish among the different p53 variants; mutant $\text{p53}^{\text{R175H}}$ aggregates had a wider dispersion of fluorescent spot intensities compared to wildtype p53, which likewise had a wider dispersion than mutant $\text{p53}^{\text{L344P}}$ monomers. We quantified the dispersion of these intensity distributions using the Fano factor, defined as the variance in intensity of an image divided by the mean intensity. The Fano factor serves as an index of aggregation status. For instance, $\text{p53}^{\text{R175H}}$ complexes showed a higher Fano factor than $\text{p53}^{\text{L344P}}$ monomers. There was a linear relationship between Fano factor and spot number for all p53 conformational variants, each with distinct slopes. Therefore, by plotting standard curves for Fano factor versus spot number for each of the conformational variants, the aggregation status of p53 could be determined.

Single-molecule anti-p53 autoantibody detection

Recombinant human p53 protein was biotinylated with EZ-Link Sulfo-NHS-Biotin reagent and purified by gel filtration chromatography on 7 kD columns as described above. Biotinylated protein was stored at -20°C in PBS containing 20% glycerol and 0.1% sodium azide. The purified biotinylated protein was coated in one channel of a dual-channel single-molecule microfluidic capture chip via a streptavidin linker (Thermo Fisher Scientific); the other channel was kept uncoated as a background control. Streptavidin was used instead of NeutrAvidin (a deglycosylated form of avidin protein found in chicken egg white) since most individuals had large amounts of circulating anti-NeutrAvidin IgG, probably because these people eat eggs. Human plasma samples were diluted $10^3$-$10^6$× in SMAC$^{BSA}$ buffer and passed continuously through both channels of the SMAC chip for 2 hr by oscillating flow. The chip was then washed with SMAC$^{BSA}$ buffer (1 ml) and incubated with Alexa Fluor 488-labeled goat anti-human IgG (1-10 nM) for 30 min. The chip was washed again and visualized by TIRF microscopy. The spots in the uncoated channel reflected the amount of basal human IgG deposited on the chip via non-specific binding. Autoantibody levels were hence calculated by subtracting this number of non-specific IgG counts from total counts in the p53-coated channel.

<u>qPCR</u>

Serum DNA was extracted with the Plasma/Serum Cell-Free Circulating DNA Purification Micro Kit (Norgen Biotek) according to manufacturer instructions. Briefly, 50 µl of mouse serum was collected and eluted with nuclease-free water (50 µl). The eluate (2 µl) was then used for PCR amplification with 2× SsoFast EvaGreen Supermix (Bio-Rad) (10 µl), 5 µM GFP primer mix (2 µl), and nuclease-free water (6 µl) on a CFX96 qPCR system (Bio-Rad) with the following thermal cycling conditions: 98°C for 1 min followed by 60 cycles of 98°C for 5 sec and 60°C for 10 sec. A melt curve was performed from 65°C to 95°C. To generate qPCR standard curves for *gfp* and *p53*, *gfp* and *hp53* plasmid DNA (2 pg), respectively, were serially diluted. Standard curves displayed cycle threshold ($C_t$) values as a function of DNA copy number. Primer pairs for p53 qPCR were: 5' CCTTGCCGTCCCAAGCA 3' (forward) and 5' GTGTAGGAGCTGCTGGTG 3' (reverse). Primer pairs for GFP qPCR were: 5' ACGTAAACGGCCACAAGTTC 3' (forward) and 5' AAGTCGTGCTGCTTCATGTG 3' (reverse).

<u>p53 native protein gel electrophoresis</u>

BHK21 cells were transfected using Lipofectamine 2000 with mutant or wildtype p53 constructs (0.1-20 µg DNA) in 6-well plates. After 16 hr, lysate was prepared as described above with 18 mM CHAPS in TBS containing DNase and protease inhibitor. Lysate was added with 20% glycerol and 5 mM Coomassie G-250 dye then loaded onto a 3-12% native PAGE Bis-Tris gel (Invitrogen). Electrophoresis was performed in 50 mM Bis-Tris and 50 mM Tricine plus 0.02% Coomassie G-250 dye in the cathode buffer for 2 hr at 100 V. Proteins were transferred to a polyvinylidene membrane and stained with Coomassie G-250 dye. The membrane was fixed with 8% acetic acid for 20 min and destained with 100% methanol. p53 proteins were detected by immunoblot with DO-1 antibodies and HRP-conjugated anti-mouse secondary antibodies.

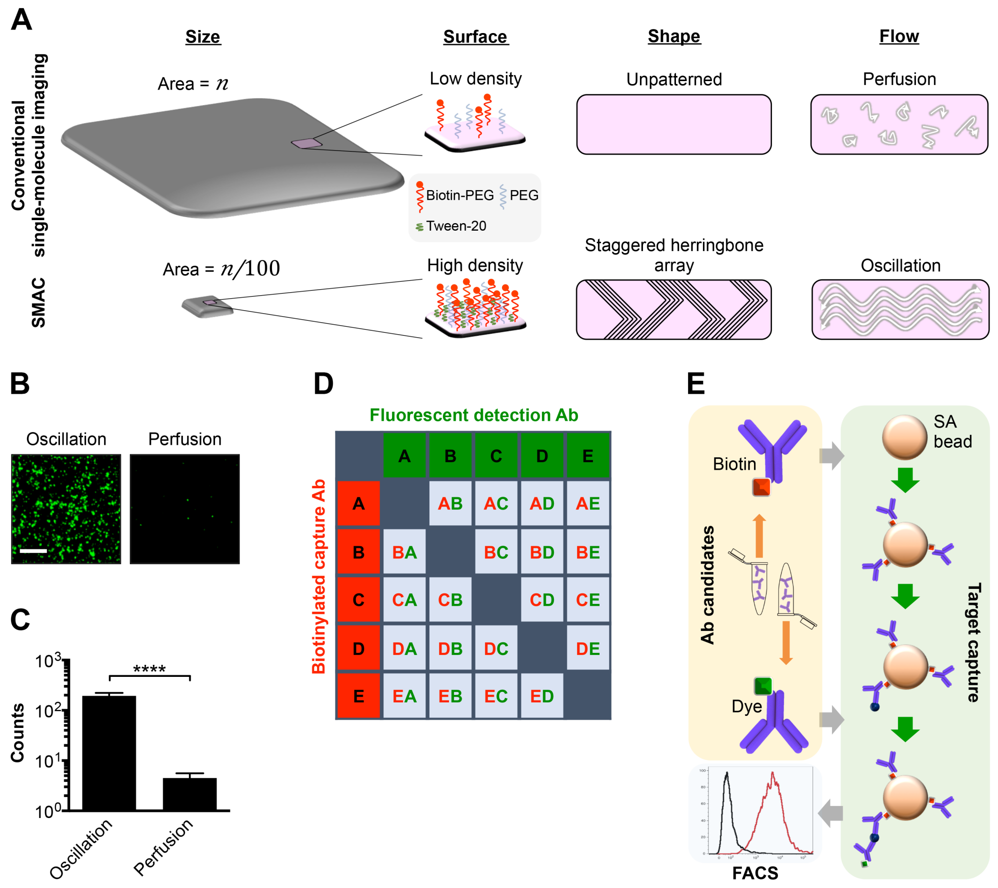


**Fig. S1. SMAC chip design and target protein capture platform.** (**A**) Schematic diagram depicting features of SMAC (bottom), contrasted to conventional single-molecule imaging methods (top), that enable single-molecule imaging of blood samples at sub-femtomolar sensitivity. The miniature size, high-density capture surface, patterned channel shape, and continuous oscillating flow scheme of the SMAC chip synergize to efficiently concentrate proteins-of-interest on the chip. **(B, C)** SMAC images (**B**) and quantification (**C**) of purified GFP pulled down on the SMAC chip via either oscillating flow or perfusion in aqueous buffer. (**D**, **E**) Bead-based flow cytometry assay for screening capture and detection antibody (Ab) pairs. Schematic of possible capture/detection reagent combinations (**D**) and the assay setup (**E**). In (**E**), biotinylated capture antibody was loaded onto streptavidin (SA)-coated magnetic beads and incubated with dye-labeled detection antibody in the presence or absence of the corresponding target protein. The beads were then washed and examined by flow cytometry (FACS) to determine an index of the binding affinity of capture/detection antibody pairs. Data are expressed as mean ± SD. *****P*<0.0001. *P*-values are from a two-sided unpaired *t*-test. Scale bar, 4 µm.

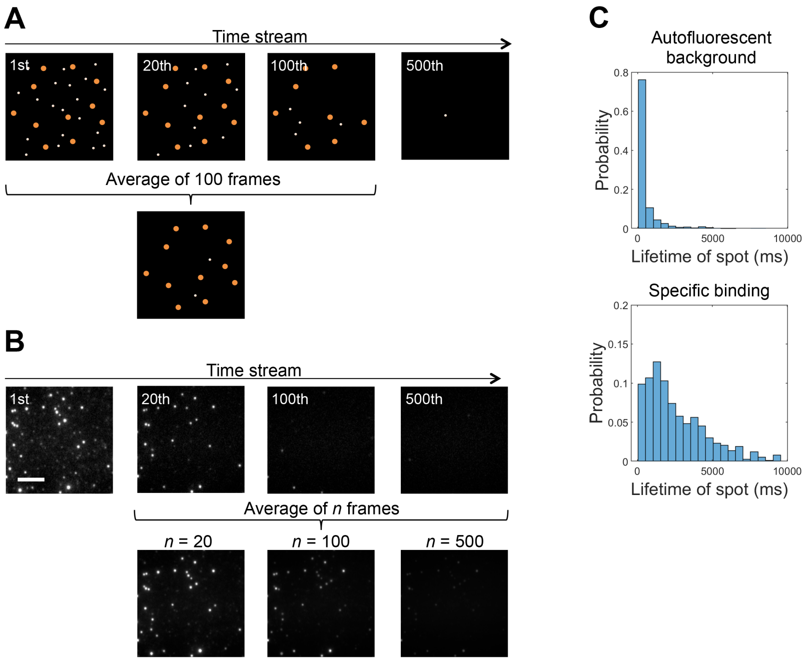


**Fig. S2. Single-molecule imaging and analysis components of SMAC.** Illustration of TIRF time-stream processing to average out diffusive background and non-specific binding signals. Schematic (**A**) and actual empirical data (**B**) depicting outcomes of this procedure using different time-average windows (*n*). In (**A**), orange and white spots indicate specific antibody clusters and non-specific antibody, respectively. (**C**) Comparison of the fluorescent spot lifetime of autofluorescent background versus specific binding under time-stream imaging with continuous laser excitation. Note: the average lifetime of autofluorescent background is $585 \pm 1.38\times10^3$ ms, while the lifetime of specific binding is $2.9\times10^3 \pm 2.87\times10^3$ ms.

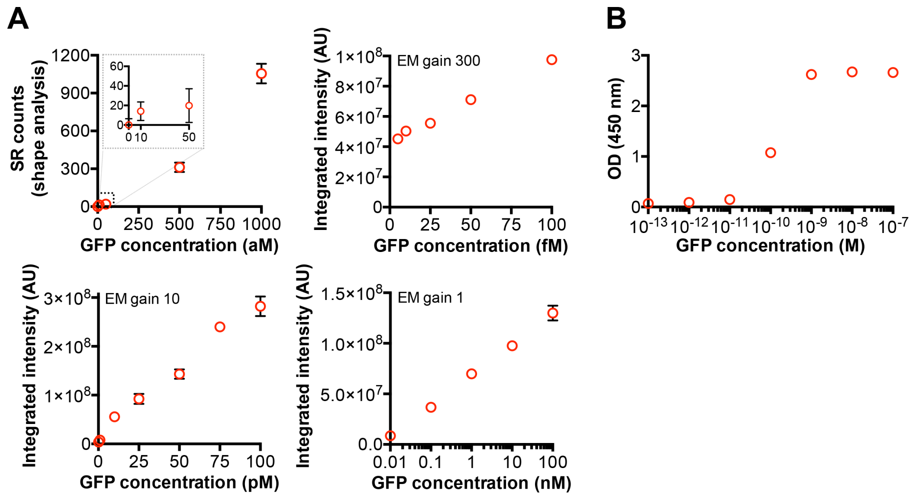


**Fig. S3. Comparison of GFP detection by SMAC or ELISA.** (**A**) Purified GFP at different concentrations (10 aM to 100 nM) was added into aqueous buffer and examined by SMAC with either shape analysis (top left, EM gain 300) or intensity analysis at different EM gain settings. SMAC detection of GFP had sub-fM sensitivity and a dynamic range from <1 fM to 100 nM. (**B**) ELISA detection of purified GFP. ELISA had a sensitivity of ~10 pM and a dynamic range from 10 pM to 1 nM.

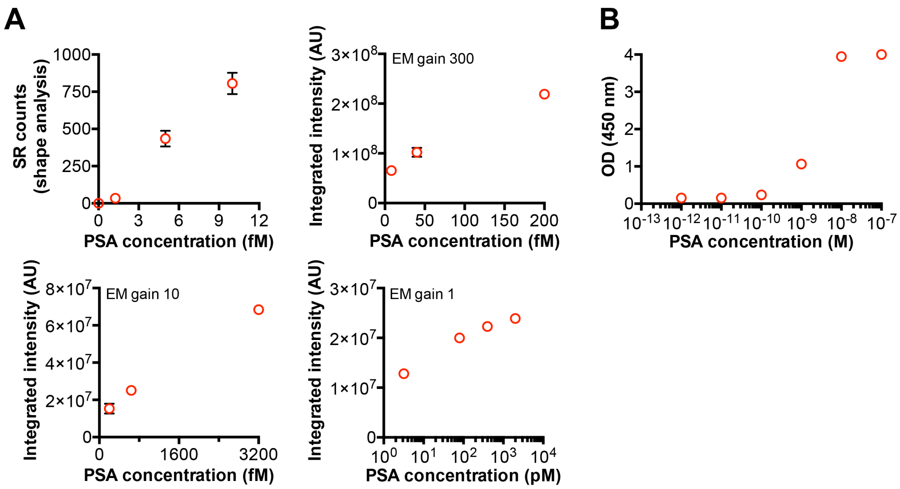


**Fig. S4. Comparison of PSA detection by SMAC or ELISA.** (**A**) Purified PSA at different concentrations (1 fM to 2 nM) was added into aqueous buffer and examined by SMAC with either shape analysis (top left, EM gain 300) or intensity analysis at different EM gain settings. SMAC detection of PSA had a sensitivity limit of <1 fM and a dynamic range from 1 fM to 1 nM. (**B**) ELISA detection of purified PSA. ELISA had a sensitivity limit of ~10 pM and was a dynamic range from 10 pM to 1 nM.

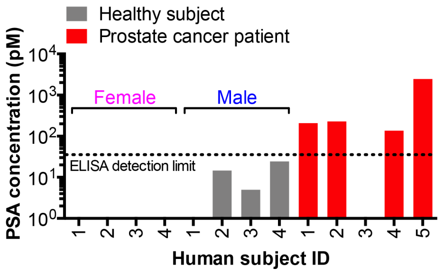


**Fig. S5. ELISA of PSA levels in plasma from prostate cancer patients or healthy donors.** PSA protein levels were measured by ELISA in plasma collected from prostate cancer patients (*n* = 5; red bars), as well as healthy male (*n* = 4; gray bars) or female (*n* = 4; gray bars) blood donors (all subjects from the same cohort as in Fig. 2G and Table S1). Note: 50 µl of plasma was required to detect PSA from prostate cancer patients (compared to 4 µl used for SMAC), and basal circulating PSA levels in healthy male blood donors fell below the ELISA detection limit (dotted line).

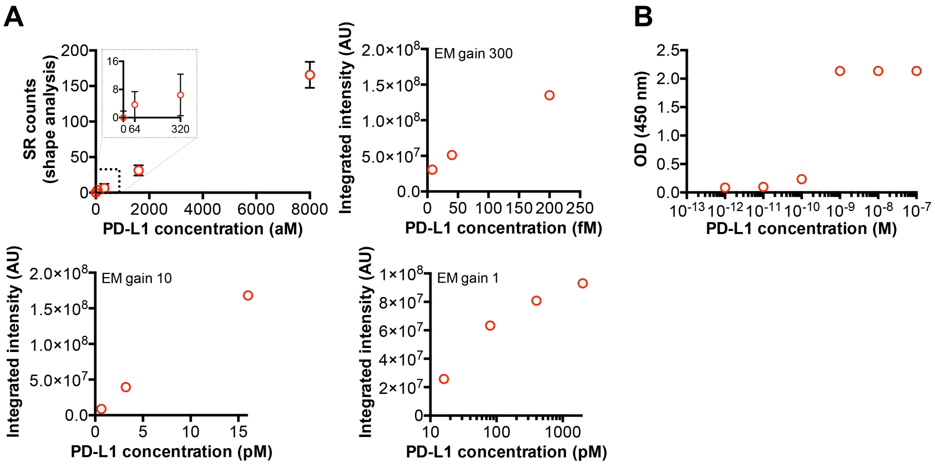


**Fig. S6. Comparison of PD-L1 detection by SMAC or ELISA.** (**A**) Purified PD-L1 at different concentrations (64 aM to 2 nM) was added into aqueous buffer and examined by SMAC with either shape analysis (top left, EM gain 300) or intensity analysis at different EM gain settings. SMAC detection of PD-L1 had a sensitivity limit of <1 fM and a dynamic range from <1 fM to 1 nM. (**B**) ELISA detection of purified PSA. ELISA had a sensitivity limit of ~10 pM and a dynamic range from 10 pM to 1 nM.

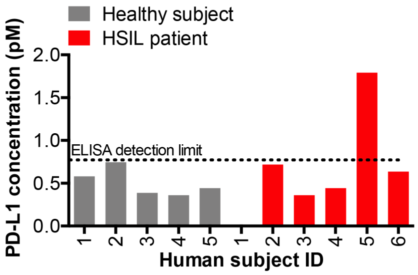


**Fig. S7. ELISA of PD-L1 levels in plasma from high-grade cervical squamous epithelial neoplasia (HSIL) patients or healthy female blood donors.** PD-L1 protein levels were measured by ELISA in plasma collected from HSIL patients ($n$ = 6; red bars), as well as healthy female ($n$ = 5; gray bars) blood donors (all subjects from the same cohort as in Fig. 2K and Table S2). Note: basal circulating PD-L1 levels in healthy blood donors fell below the ELISA detection limit (dotted line).

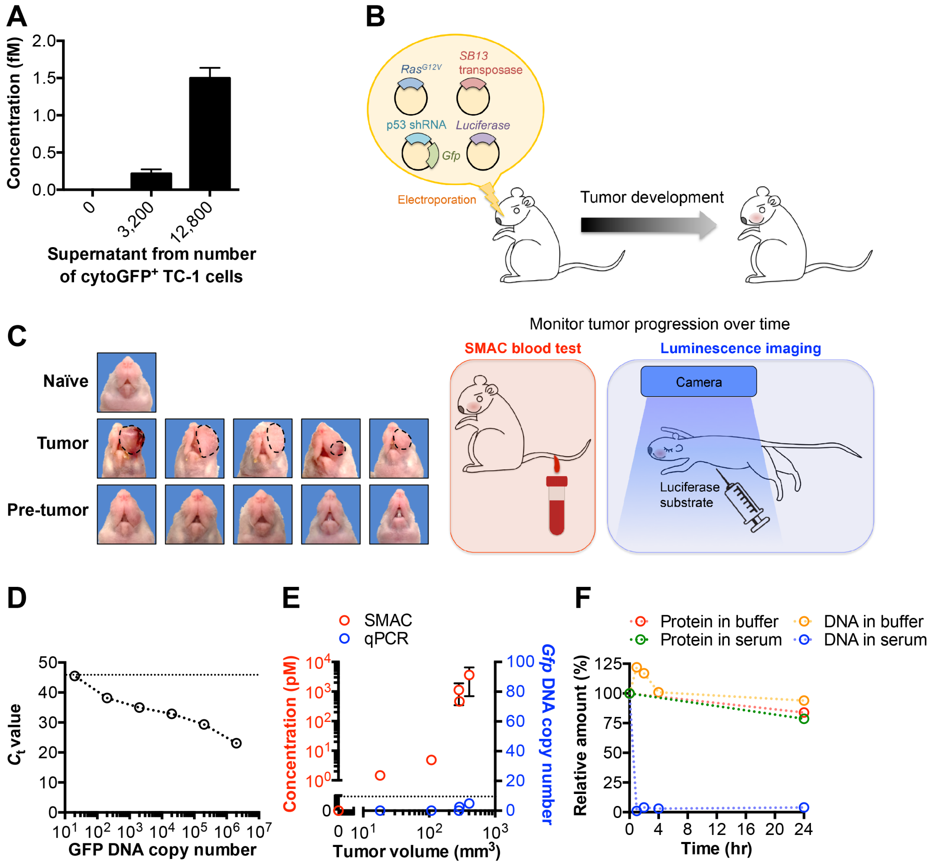


**Fig. S8. Comparison of tumor intracellular protein versus DNA release into serum of mice by SMAC and qPCR, respectively.** (**A**) SMAC quantification of cytoGFP released from a cultured cytoGFP$^+$ tumor cell line (TC-1) into the supernatant. (**B**) Schematic diagram of the preclinical tumor model setup and design of the SMAC and luminescence imaging experiments. (**C**) Photographs of mice not induced with tumor (labeled 'naïve'), as well as mice induced with tumor that either displayed grossly visible tumor after more than two months (labeled 'tumor') or had no signs of tumor (labeled 'pre-tumor'). SMAC and luminescence results are shown in Fig. 3C. (**D**) Standard curve depicting the performance of qPCR for GFP DNA detection over a range of 20 to $2 \times 10^6$ DNA copies (goodness of fit $R^2 = 0.9626$). The dotted line indicates assay background. (**E**) Serum cytoGFP protein levels (measured by SMAC; red circles) versus *cyto-gfp* DNA levels (measured by qPCR; blue circles) in mice with visible tumor ($n = 5$; mice pictured in (**C**) as a function of tumor volume. Circulating cytoGFP levels increased exponentially with transplanted tumor growth and ranged from 1 pM to 1 nM. The dotted line indicates the detection limit of qPCR. (**F**) Comparison of GFP protein versus DNA stability over time at 37°C in mouse serum or aqueous buffer, measured by SMAC and qPCR, respectively. An identical molar amount of protein and DNA was added in each group. All data are expressed as mean ± SD.

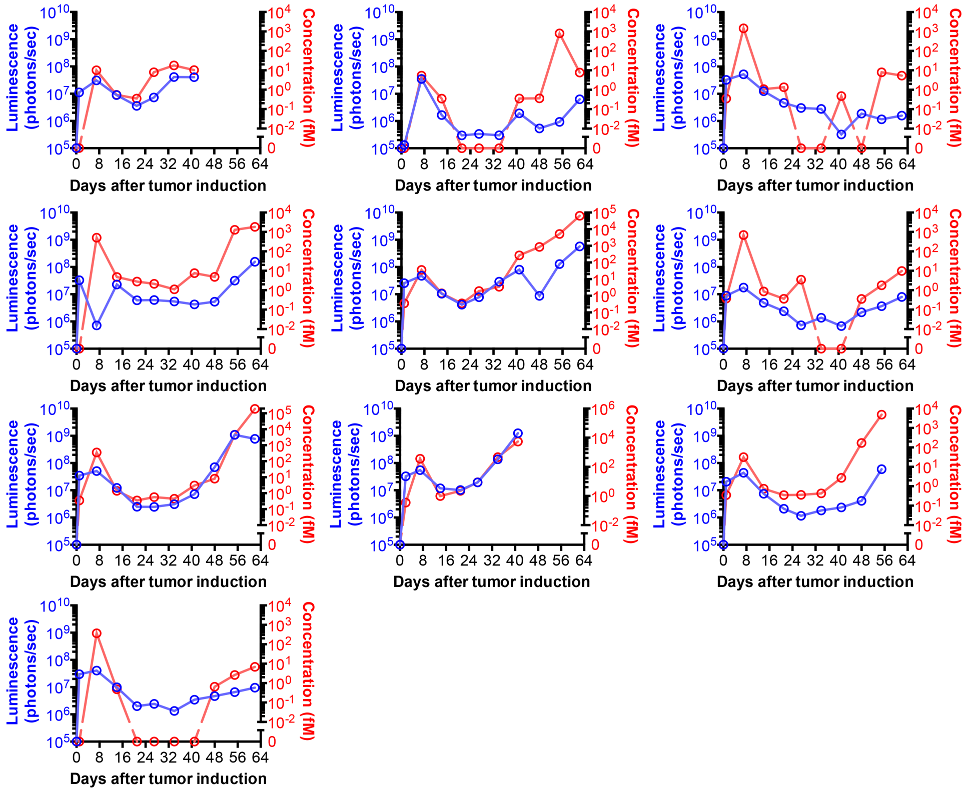


**Fig. S9. Temporal progression of tumor burden and circulating intracellular protein levels in mice induced with spontaneous tumor.** Spontaneous tumor formation in mice ($n$ = 10; same cohort as in Fig. 3D, E) was induced via intrabuccal delivery of *Ras*$^{G12V}$, *shP53, cyto-gfp*, and *luciferase* DNA (schematic in Fig. S8B). At different time points over the first two months, serum was collected, and circulating cytoplasmic GFP levels (red circles) were measured by SMAC. Circulating cytoGFP levels ranged from 1 fM to 100 pM throughout this experiment. Buccal luciferase activity (blue circles) was measured by luminescence imaging as an index of tumor burden.

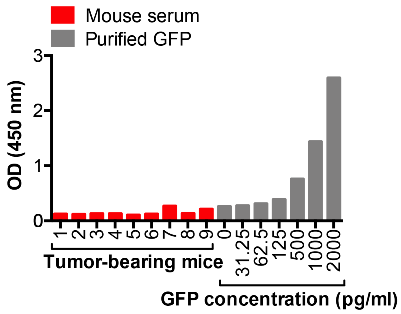


**Fig. S10. ELISA of intracellular protein levels in the blood in tumor-bearing mice.** GFP protein levels (red bars) were measured by ELISA in serum collected from tumor-bearing mice (*n* = 9; same cohort as in Fig. 3D, e and Fig. S9) more than two months after tumor was induced. One of the mice had died by this time and was thus not included. For comparison, an ELISA standard curve with purified GFP is shown (gray bars).

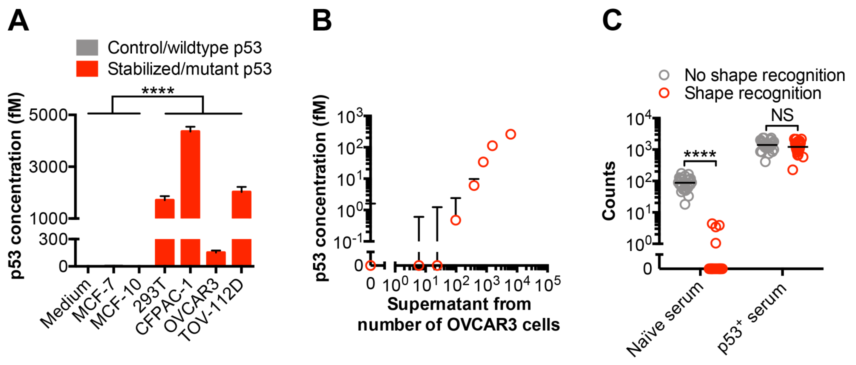


**Fig. S11. Detection of mutant protein release from cancer cells by single-molecule imaging.** (**A**) SMAC with shape analysis of p53 levels in lysate of a panel of cell lines carrying mutant p53 (CFPAC-1 (C242R), OVCAR3 (R248Q), TOV-112D (R175H)) or wildtype p53 (MCF-7, MCF-10). A total of 1,000 cells was used for each group. (**B**) Shape analysis of mutant $p53^{R248Q}$ levels in 0.2 μm-filtered supernatant from a human ovarian cancer cell line (OVCAR3) cultured overnight. (**C**) Comparison of human p53 SMAC results without (gray circles) or with (red circles) error correction using shape analysis. In these experiments, naïve mouse serum and serum spiked with human p53 (1 pM) were run over multiple independent trials ($n$ = 26). Data in (**A**, **B**) are expressed as mean ± SD; individual data points with mean are shown in (**C**). ****$P$<0.0001; NS, not significant. $P$-values are from a two-sided unpaired $t$-test.

**A**

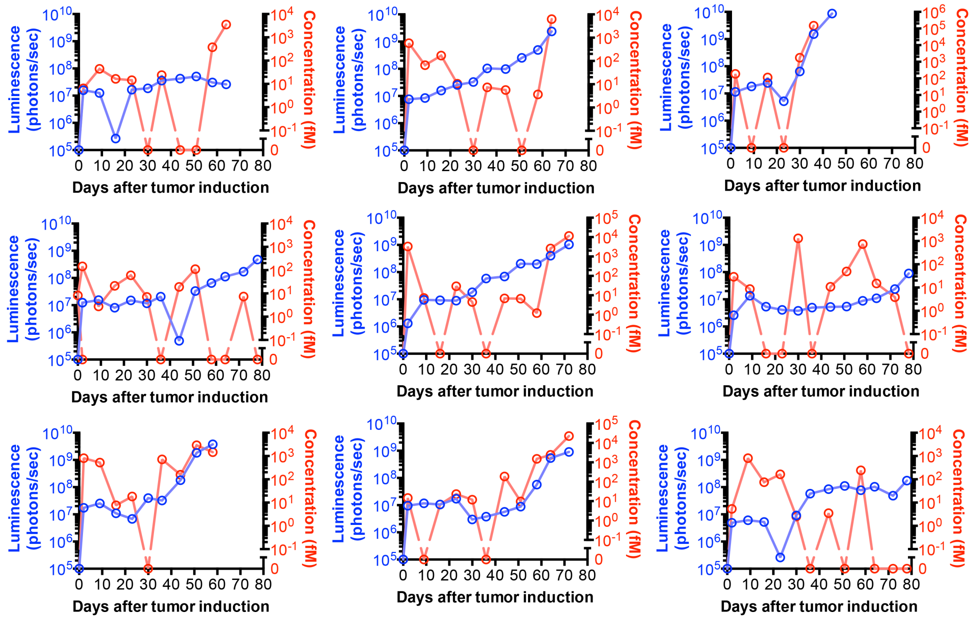


**B**

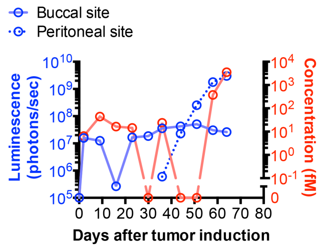


**Fig. S12. Temporal progression of tumor burden and circulating mutant protein levels in mice induced with spontaneous tumor.** (**A**) Spontaneous tumor formation in immune-deficient mice ($n$ = 10; same cohort as in Fig. 3H, I) was induced via delivery of human *p53*$^{R175H}$, *Ras*$^{G12V}$, and *luciferase* DNA to the buccal mucosa. At different time points over the first two months, serum was collected, and circulating mutant p53 levels (red circles) were measured by SMAC. Local buccal luciferase activity (blue circles) was measured by luminescence imaging as an index of tumor burden. Circulating p53 concentrations ranged from 10 fM to 1 nM and correlated with tumor stage. (**B**) In one of these mice, peritoneal metastasis of the buccal tumor was observed. Circulating mutant p53 levels in this mouse, as assessed by SMAC, paralleled the rise in peritoneal luminescence.

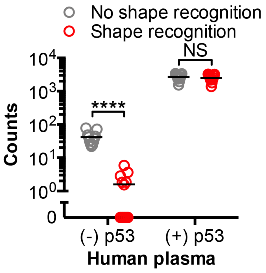


**Fig. S13. Error correction outcomes using single-molecule shape analysis for rare proteins in human blood.** Comparison of human p53 SMAC results without (gray circles) or with (red circles) error correction using shape analysis. In these experiments, plasma from individual healthy blood donors ($n$ = 10) was either spiked with human p53 (2 pM) or left untreated and then examined by SMAC. Individual data points with mean are shown. ****$P$<0.0001; NS, not significant. $P$-values are from a two-sided unpaired $t$-test.

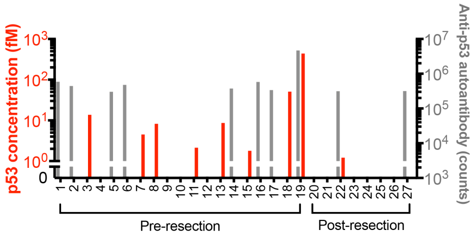


**Fig. S14. Quantification of circulating mutant p53 and endogenous anti-p53 autoantibodies from ovarian cancer patients with stage III disease either before or after surgical resection.** Data for individual human plasma samples are expressed as mean ± SE.

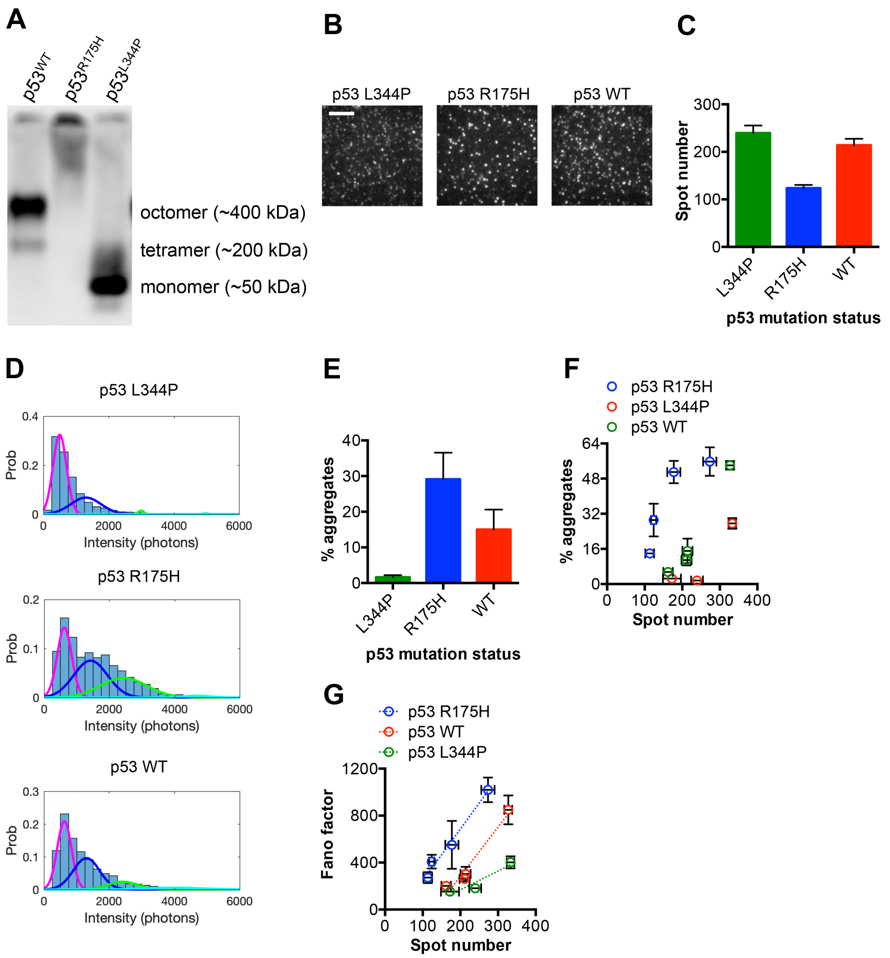


**Fig. S15. Aggregation status of wildtype or mutant p53 conformational variants.** (**A**) Native PAGE and immunoblot of mutant and wildtype (wt) p53 to identify their oligomerization states. BHK21 cells were transfected with *p53*$^{WT}$, *p53*$^{R175H}$, or *p53*$^{L344P}$ cDNA linked to a GFP expression cassette. (**B**, **C**) TIRF microscope images (**B**) and quantification (**C**) of fluorescent spots from equal total concentrations of individual mutant or wildtype p53 proteins and complexes. (**D**, **E**) Relationship between aggregate formation and p53 mutation status. (**D**) Gaussian fitting of intensity histograms for p53$^{L344P}$, wildtype p53, and p53$^{R175H}$. The curves show the intensity distributions for monomer (pink), dimer (blue), or tetramer (green), and aggregate (aqua) populations of p53. (**E**) Percentage of aggregates in each group, calculated as the combined area under the tetramer and aggregate Gaussian curves in (**D**) divided by the total histogram area. For these experiments, identical concentrations were used for each sample. (**F**) Relationship between spot number and percentage of aggregates for different p53 conformational variants across a range of concentrations. (**G**) Aggregation index of the various wildtype and p53 mutants, expressed as Fano factor to reflect the width of the intensity distribution of p53 in each sample (see Methods for calculation details).

| ID | Age | Sex | Race | Tumor type |
|---|---|---|---|---|
| **1** | 61 | M | Caucasian | Adenocarcinoma of the prostate gland |
| **2** | 65 | M | Caucasian | Adenocarcinoma of the prostate gland |
| **3** | 72 | M | Caucasian | Adenocarcinoma of the prostate gland |
| **4** | 64 | M | Caucasian | Adenocarcinoma of the prostate gland |
| **5** | 58 | M | Caucasian | Adenocarcinoma of the prostate gland |

**Table S1. Clinical characteristics of prostate cancer patient samples**

| ID | Age | Sex | Race | Tumor type |
|---|---|---|---|---|
| **1** | 20 | F | Caucasian | High-grade squamous intraepithelial lesion (CIN 2) |
| **2** | 26 | F | Caucasian | High-grade squamous intraepithelial lesion |
| **3** | 69 | F | African American | High-grade squamous intraepithelial lesion |
| **4** | 38 | F | Caucasian | High-grade squamous intraepithelial lesion (CIN 3) |
| **5** | 51 | F | African American | High-grade squamous intraepithelial lesion (CIN 2) |
| **6** | 65 | F | Caucasian | High-grade squamous intraepithelial lesion (CIN 3) |

**Table S2. Clinical characteristics of HPV-induced cervical intraepithelial lesion patient samples**

| ID | Age | Sex | Race | Tumor type* | FIGO stage |
|---|---|---|---|---|---|
| **1** | 77 | F | Caucasian | Serous carcinoma | pT3 |
| **2** | 32 | F | Caucasian | Metastatic clear cell carcinoma | pT4 |
| **3** | 55 | F | Caucasian | Metastatic serous carcinoma | N/A |
| **4** | 74 | F | Caucasian | Metastatic serous carcinoma | pT3c |
| **5** | 59 | F | Caucasian | Serous carcinoma | pT3c |
| **6** | 75 | F | European | Serous adenocarcinoma | pT4 |
| **7** | 49 | F | Caucasian | Metastatic serous carcinoma | pT3c |
| **8** | 88 | F | Caucasian | Metastatic serous carcinoma | pT3c |
| **9** | 60 | F | African American | Adenocarcinoma | pT3c |
| **10** | 64 | F | Caucasian | Metastatic serous adenocarcinoma | N/A |
| **11** | 64 | F | Caucasian | Serous adenocarcinoma | N/A |
| **12** | 68 | F | Caucasian | Serous carcinoma | pT3c |
| **13** | 64 | F | Caucasian | Metastatic serous carcinoma | pT3c |
| **14** | 61 | F | Caucasian | Metastatic serous carcinoma | pT4 |

* All tumors were of ovarian origin and classified as high-grade. N/A: not available.

**Table S3. Clinical characteristics of ovarian cancer patient samples**

| ID | Tumor type | Tumor grade | Prior resection | FIGO stage | Mutation |
|---|---|---|---|---|---|
| **1** | Serous carcinoma | High-grade | No | pT3cN1MX | TP53 p.S127fs |
| **2** | Serous carcinoma | High-grade | No | pT3aN1MX | TP53 p.Y163C |
| **3** | Serous carcinoma | High-grade | No | pT3cN1MX | TP53 p.Y163C |
| **4** | Undifferentiated carcinoma | High-grade | No | pT3cN1MX | TP53 g.7576852C>T (splice site) |
| **5** | Clear cell carcinoma | -- | No | pT3bN0MX | TP53 p.F341fs |
| **6** | Serous carcinoma | High-grade | No | pT3cN1MX | TP53 p.P278S |
| **7** | Serous carcinoma | High-grade | No | pT3cN0MX | TP53 p.E366fs |
| **8** | Serous carcinoma | High-grade | No | pT3cNXMX | TP53 p.T211A |
| **9** | Carcinosarcoma | -- | No | pT4bN0MX | TP53 p.V218G |
| **10** | Serous carcinoma | High-grade | No | pT3cN1MX | TP53 p.R306* |
| **11** | Poorly differentiated carcinoma | High-grade | No | pT3cN1MX | TP53 p.A353V |
| **12** | Serous carcinoma | High-grade | No | pT3NXMX | TP53 p.R267P |
| **13** | Poorly differentiated carcinoma | High-grade | No | pT3cN0MX | TP53 p.F341fs |
| **14** | Serous carcinoma | High-grade | No | pT3cN0M0 | TP53 p.L264fs |
| **15** | Serous carcinoma | High-grade | No | pT3cN1MX | TP53.S127F |
| **16** | Adenocarcinoma | High-grade | No | pT3cN0MX | TP53 p.P177R |
| **17** | Serous carcinoma | High-grade | No | pT3cN1M0 | TP53 p.R249G |
| **18** | Serous carcinoma | High-grade | No | pT3cN1MX | TP53 p.R273L |
| **19** | Serous carcinoma | High-grade | No | pT3cN1MX | TP53 p.S241F |
| **20** | Serous carcinoma | High-grade | Yes | pT3 | TP53 p.D259V |
| **21** | Serous carcinoma | High-grade | Yes | pT3 | TP53 p.S241F |
| **22** | Serous carcinoma | High-grade | Yes | pT3 | TP53 p.H168Q |
| **23** | Serous carcinoma | High-grade | Yes | pT3 | TP53 p.R175H |
| **24** | Adenocarcinoma | High-grade | Yes | pT3 | TP53 p.T155N |
| **25** | Adenocarcinoma | High-grade | Yes | pT3 | TP53 p.M340fs |
| **26** | Serous carcinoma | High-grade | Yes | pT3 | TP53 g.7579311C>G (splice site) |
| **27** | Adenocarcinoma | High-grade | Yes | pT3 | TP53 g.7576927C>T (splice site) |

**Table S4. Pathologic features of tumors in late-stage ovarian cancer patients**

| ID | Tumor type | Tumor grade | Prior resection | FIGO stage | Mutation |
|---|---|---|---|---|---|
| **1** | Mixed adenocarcinoma | High-grade | No | pT1 | CTNNB1 p.S33Y |
| **2** | Carcinoma with mucinous features | Low-grade | No | pT1cN0MX | PIK3CA p.H1047R |
| **3** | Non-invasive micropapillary carcinoma | Low-grade | No | pT2N0MX | CDKN2A p.R87Q |
| **4** | Serous borderline tumor | -- | No | pT1aN0MX | TP53 p.V216L |
| **5** | Carcinoma (unspecified type) | -- | No | pT1 | TP53 p.K372fs |
| **6** | Clear cell carcinoma | -- | No | pT1aN0MX | TP53 p.P300fs |
| **7** | Non-invasive micropapillary carcinoma | Low-grade | No | pT1aN0MX | TP53 p.R249G |

**Table S5. Pathologic features of tumors in early-stage ovarian cancer patients**
